# Tomosyns attenuate SNARE assembly and synaptic depression by binding to VAMP2-containing template complexes

**DOI:** 10.1101/2023.06.20.545669

**Authors:** Marieke Meijer, Miriam Öttl, Jie Yang, Aygul Subkhangulova, Avinash Kumar, Alexander J. Groffen, Jan R.T. van Weering, Yongli Zhang, Matthijs Verhage

**Affiliations:** Department of Human Genetics, Center for Neurogenomics and Cognitive Research, Amsterdam University Medical Center, 1081HV Amsterdam, The Netherlands; Department of Functional Genomics, Center for Neurogenomics and Cognitive Research, Vrije Universiteit Amsterdam, 1081HV Amsterdam, The Netherlands; Department of Cell Biology, Yale School of Medicine, New Haven, CT 06511, USA; Department of Molecular Biophysics and Biochemistry, Yale University, New Haven, CT 06511, USA

**Author notes:** Correspondence (Y.Z), (M.V.). These authors contributed equally (Marieke Meijer, Miriam Öttl, Jie Yang). Co-senior authors (Yongli Zhang, Matthijs Verhage).

**Keywords:** Tomosyn, STXBP5, STXBP5L, membrane fusion, SNARE assembly, template complex, optical tweezers, synaptic strength, short-term plasticity, vesicle priming

## Abstract

Tomosyns are soluble SNARE proteins proposed to attenuate membrane fusion by competing with synaptobrevin-2/VAMP2 for SNARE-complex assembly. Here, we present evidence against this scenario using a novel mouse model, energy barrier recordings, and single-molecule force measurements. Tomosyn-1/2 deficiency drastically enhanced the probability that synaptic vesicles fuse at synapses, resulting in stronger synapses with faster depression and slower recovery. While wildtype tomosyn-1m rescued these phenotypes, substitution of its SNARE motif with that of synaptobrevin-2/VAMP2 did not. Force measurements revealed that tomosyn’s SNARE motif cannot substitute synaptobrevin-2/VAMP2 to form template complexes with Munc18-1 and syntaxin-1, an essential intermediate for SNARE assembly. Instead, tomosyns bind synaptobrevin-2/VAMP2-containing template complexes and prevent SNAP-25 association. Structure-function analyses indicate that regions outside the SNARE motif contribute to tomosyn’s inhibitory function. These results reveal that tomosyns regulate synaptic transmission by preventing SNAP-25 binding to template complexes, increasing the energy barrier for synaptic vesicle fusion, and limiting synaptic depression.

## Introduction

SNARE-dependent membrane fusion is required for the secretion of most chemical signals ^1, 2^. As minimal fusion machinery, SNAREs are conserved from yeast to mammals and couple their folding and assembly to membrane fusion ^3–5^. The fully assembled SNARE complex is a four-helix bundle formed by characteristic SNARE motifs. While SNARE complexes can spontaneously form *in vitro* with low speed and specificity, their assembly *in vivo* is controlled by many proteins, such as Munc18s, Munc13s, synaptotagmins, and complexins ^2, 6–8^. Mutations in the genes encoding SNARE and regulatory proteins lead to a wide range of human diseases, including various neurodevelopmental, cardiovascular and hematological disorders ^8–12^. However, many aspects of regulated SNARE assembly and its coupling to membrane fusion remain poorly understood.

Synaptic vesicle (SV) fusion is preceded by a dynamic multi-step process involving docking the SV onto the presynaptic membrane and priming the SV for rapid fusion upon stimulation ^13–16^. These steps are associated with distinct intermediates of assembly of three synaptic SNAREs: syntaxin-1, SNAP-25, and synaptobrevin-2(syb2)/VAMP2. The template complex constitutes a key SNARE intermediate for SNARE assembly ^17–23^. In this complex, syntaxin-1 and syb2/VAMP2 bind to the surface of Munc18-1 such that the N-terminal regions of their SNARE motifs are aligned, while their C-terminal regions are kept separated. SNAP-25 specifically and rapidly associates with the template complex, leading to full SNARE assembly ^19, 24^. Intermediates of SNARE assembly might be important targets for regulating membrane fusion, especially the fusion probability of a primed vesicle. Such regulation not only determines synaptic strength, but also set the short-term plasticity properties of synapses (e.g. facilitating/’tonic’ or depressing/’phasic’), both of which help determine the computational properties of brain circuits ^25, 26^.

Tomosyns play important roles in vesicle fusion ^27^, but their mechanism of action remains unclear. Tomosyn isoforms share a conserved structure containing a large N-terminal WD40 repeat domain and a small C-terminal SNARE motif ^28, 29^. In cell-free assays, the tomosyn SNARE motif competes with the homologous SNARE motif of syb2/VAMP2, resulting in a tomosyn SNARE complex that is expected to be non-fusogenic as tomosyns lack a transmembrane domain present in VAMP2 ^30–33^. Indeed, tomosyns inhibit exocytosis in many secretory cell types ^30, 34–37^, including neurons at neuromuscular junctions in nematode and fly ^38–41^ and mammalian central and peripheral neurons ^42–44^, with a few exceptions hinting at an additional positive role ^45–47^. Mutations in tomosyns are associated with neurodevelopmental disorders and thrombosis ^11, 48–54^, suggesting an important and conserved function in different secretory tissues. However, several observations are inconsistent with the concept of tomosyns as competitive inhibitors of syb2/VAMP2. In mouse platelets and yeast, tomosyns (-orthologs) exert positive roles in secretion ^37, 55, 56^. In addition, the N-terminal WD40 domain was found to be required for its inhibitory function ^41, 57, 58^. Furthermore, NSF/αSNAP rapidly disassemble the tomosyn SNARE complex ^31^, arguing against its proposed role in SNARE assembly. These observations challenge the concept of tomosyns as competitive inhibitors of SNARE complex formation.

This study addresses these issues by combining secretion assays in living neurons, measurements of the energy barrier for fusion and single-molecule force measurements using optical tweezers. Loss of tomosyn expression lowered the energy barrier for fusion which resulted in initially stronger but fast-depressing synapses. Expression of hybrid constructs where syb2/VAMP2 replaced corresponding domains in tomosyn failed to restore these synaptic defects. In line with these observations, single-molecule force measurements showed that the tomosyn SNARE motif failed to form a template complex with Munc18-1 and syntaxin-1, indicating that tomosyns do not compete with syb2/VAMP2 during Munc18-1-chaperoned SNARE assembly. Instead, tomosyns bind to syb2/VAMP2-containing template complexes and block integration of SNAP-25, thereby lowering the fusion rate of synaptic vesicles, and limiting synaptic depression.

## Results

### A novel conditional KO mouse for both tomosyn paralogs

Due to prior complications with sub-lethal phenotypes and bleeding risk in constitutive mutant mouse models ^44, 56^, we generated a new conditional double floxed mouse model in which both tomosyn paralogs can be inactivated by Cre-mediated recombination at lox sites flanking exon 2 of tomosyn-1 (*Stxbp5*) and exon 3 of tomosyn-2 (*Stxbp5l*) (Figure S1A). Hippocampal neurons cultured from newborn mutant mice were infected with lentiviruses expressing EGFP-tagged Cre to induce recombination (cDKO), while inactive Cre (lacking the catalytic domain) was used as control ^59^. Immunoblotting confirmed that floxed neurons infected with Cre lacked a band at the expected height of 130 kDa for tomosyn-1 (Figure S1B). The conditional inactivation of tomosyn-2 has been validated before ^47^. Depleting neurons of both tomosyns did not affect the levels of the neuronal SNARE proteins syntaxin-1 and syb2/VAMP2 but resulted in a 20% reduction of SNAP-25 levels (Figures S1B and S1C). To investigate the role of tomosyns in synapse function, single hippocampal neurons were cultured on astrocyte microdots, driving neurons to form abundant synapses onto themselves, resulting in a one-neuron circuit ^60, 61^. Immunostainings showed a punctate pattern of endogenous tomosyn-1 overlapping with synaptophysin-1 puncta, confirming the synaptic enrichment of tomosyns ^40, 41, 45, 62, 63^ (Figure S1D). In cDKO neurons, the intensity of the tomosyn staining was drastically reduced (Figures S1D and S1E). Knockdown of tomosyn-1 in cultured mouse neurons has recently been reported to reduce dendritic arborization ^64^. We quantified neuronal morphology in our single-cell assay using the automated image analysis routine SynD ^65^. Two-week old autaptic cDKO neurons displayed normal dendritic length and complexity as well as synapse density (Figures S1F-H). This was also the case at an earlier time point during development (Figures S1I-L). Hence, autaptic hippocampal neurons showed no evidence of altered dendritic branching and synapse formation in the absence of both tomosyns.

### Tomosyn cDKO synapses are stronger and depress faster

Next, we performed whole-cell patch-clamp recordings on autaptic hippocampal neurons to assess the role of tomosyns in synaptic transmission. cDKO neurons displayed a 5-fold increase in the frequency of spontaneous vesicle release events (miniature excitatory postsynaptic currents, mEPSCs), while the amplitude of these events was normal (Figures 1A-C). The amplitudes of evoked EPSCs were larger in cDKO neurons (Figures 1D and 1E), while their kinetics were normal (Figure S2A-C). We performed paired-pulse recordings and calculated the paired-pulse ratio (PPR), which inversely correlates with the initial synaptic release probability (Pr) ^66, 67^. cDKO neurons showed reduced PPRs at all intervals tested (Figures 1F, 1G and S2D). Moreover, cDKO neurons displayed pronounced depression during short trains of action potentials, while control neurons largely maintained their synaptic strength (Figures 1H-K and S2E, S2F). Thus, tomosyns lower the release probability of synapses, thereby reducing their initial strength and limiting synaptic depression.

**Figure 1:**
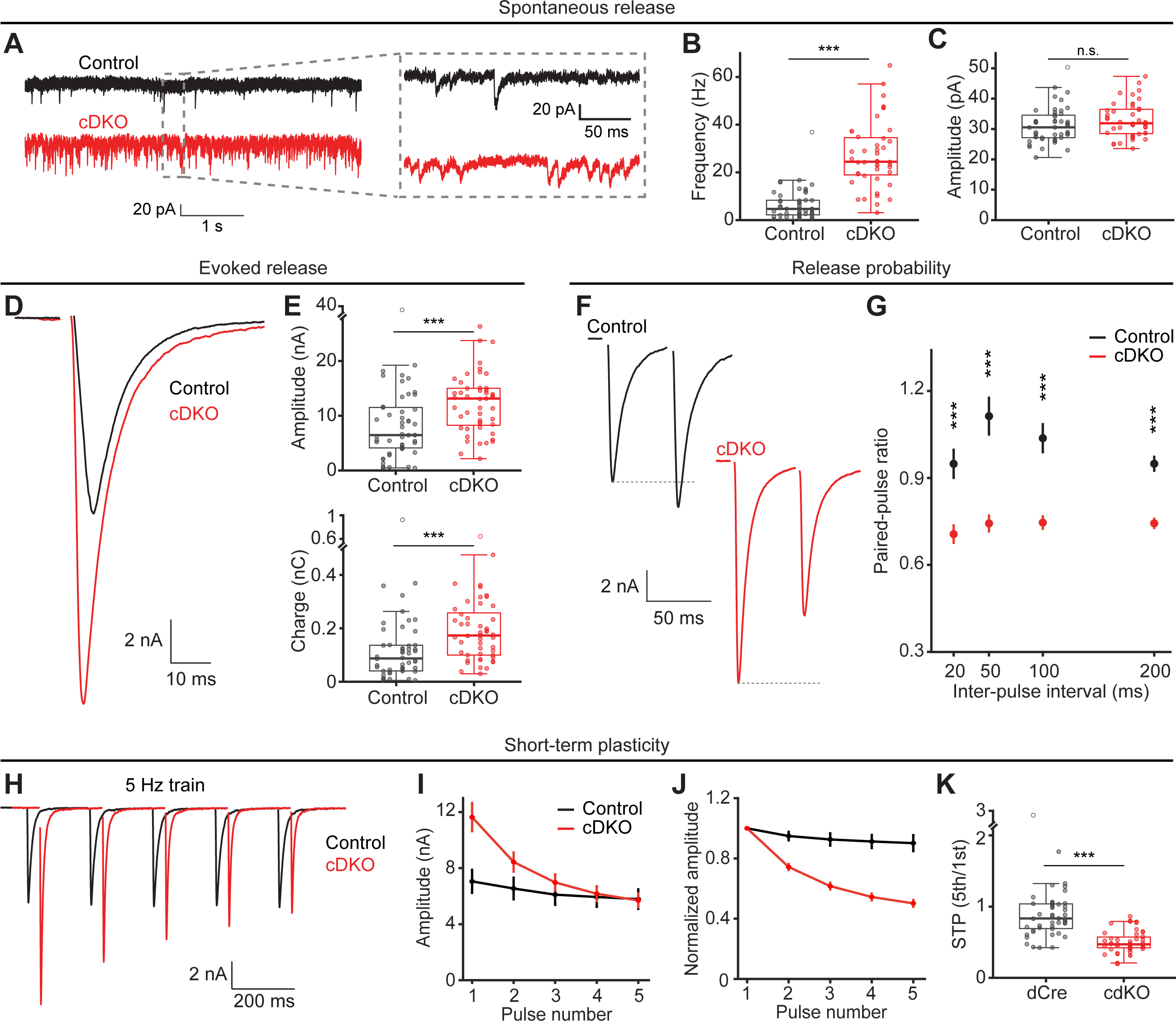
Tomosyn cDKO synapses are stronger and depress faster. (**A-C**) Analysis of spontaneous vesicle release in autaptic hippocampal neurons. Control n = 47/6, cDKO n = 44/6. (**A**) Example traces of miniature EPSCs (mEPSCs). (**B**) The frequency of mEPSCs. (**C**) mEPSC amplitude. (**D-E**) Analysis of evoked synaptic transmission. Control n = 50/6, cDKO n = 49/6. (**D**) Example traces of evoked EPSCs. (**E**, top) Amplitudes and (**E**, bottom) charges of single EPSCs. See Figure S2 A-C for quantifications of EPSC kinetics. (**F-G**) Release probability was tested by applying paired pulses with multiple inter-pulse intervals (IPIs). cDKO n = 42 – 47/6, control n = 37 – 47/6. (**F**) Example traces of paired pulses stimulated with a 50 ms IPI. (**G**) The paired-pulse ratio (PPR) was calculated by dividing the amplitude of the second pulse by the amplitude of the first pulse. See Figure S2 D for boxplots per IPI. (**H-K**) Short-term plasticity (STP) was analyzed by stimulation of 5 consecutive pulses at 5 Hz. Control n = 47/6, cDKO n = 47/6. (**H**) Example traces. (**I**) Absolute EPSC amplitudes over the 5 pulses. (**J**) STP illustrated by normalizing amplitudes to the first pulse. (**K**) STP quantified by the ratio of the fifth pulse over the first pulse. In **G**, **I** and **J**, data are presented as mean ± SEM. ***p<.001. Abbreviations: n.s. (not significant). See also Figures S1, S2 and S3.

In chromaffin cells, tomosyn-1 overexpression shifts the calcium-dependence of exocytosis ^35^. To test the calcium-dependence of synaptic responses, we applied paired pulses across a range of external calcium concentrations ([Ca^2+^]_e_) (Figure S3A). The initial EPSC size and PPR scaled with [Ca^2+^]_e_, as expected. At each concentration, PPRs were lower in cDKO neurons compared to control neurons (Figure S3B). EPSC amplitudes were normalized to flanking responses in standard 2 mM [Ca^2+^]_e_ and fitted with a Hill function (Figure S3C). The apparent calcium affinity was found to be higher (i.e., decreased *K*d for calcium) in cDKO neurons (*K*d = 1.3 ± 0.1 mM) compared to control neurons (*K*d = 1.7 ± 0.1 mM) (Figure S3D, left). The Hill coefficient, a measure of calcium cooperativity, was unchanged (Figure S3D, right). These results suggest that tomosyns reduce the Ca^2+^-sensitivity of synaptic responses.

### Tomosyns increase the energy barrier for vesicle fusion

Changes in Pr and the apparent Ca^2+^-sensitivity can result from a change in the Ca^2+^-influx upon stimulation or Ca^2+^-binding properties of the Ca^2+^-sensor for fusion, but could also be caused by a change in the intrinsic release probability of synaptic vesicles ^68, 69^. Hypertonic sucrose solutions are used to release SVs from the readily releasable pool (RRP) independent of Ca^2+^ and action potentials ^70^. Responses to 500 mM sucrose, which releases the entire RRP, were similar in tomosyn cDKO neurons and control neurons (Figures 2A and B). However, the fraction of the RRP released by a single EPSC, a common measure of the vesicular release probability (P_ves_), was increased in cDKO neurons (Figure 2C). An even stronger increase was seen when calculating the fraction of the RRP released by 250 mM sucrose (Figure 2D), which probes the intrinsic fusion efficiency, or fusogenicity, of SVs in a calcium-independent manner ^71, 72^. Together, these findings indicate that tomosyns reduce the intrinsic fusogenicity of SVs without affecting the size of the primed SV pool.

**Figure 2:**
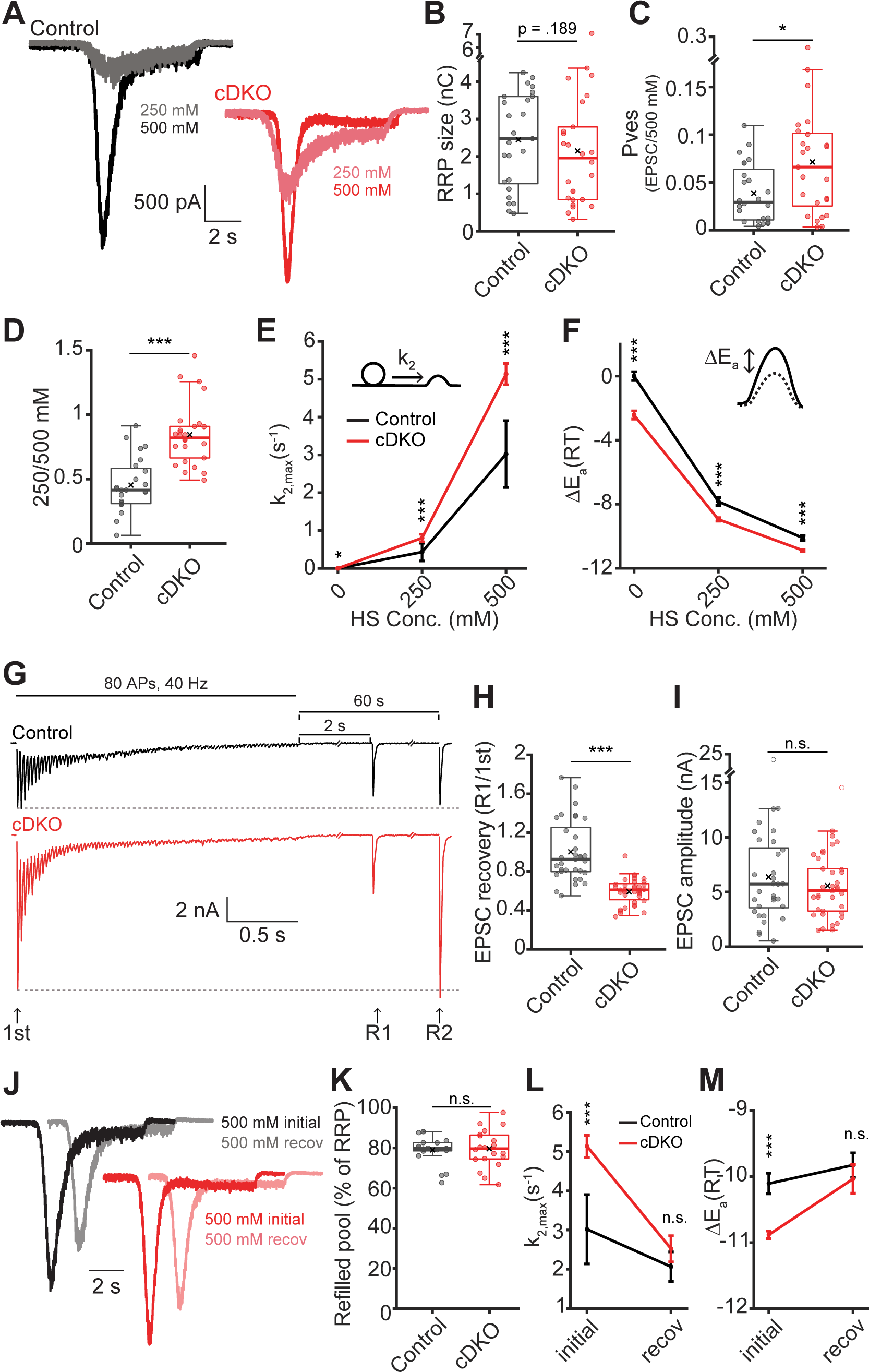
Tomosyns increase the energy barrier for vesicle fusion. (**A**) Example traces of 250 mM and 500 mM sucrose applications. (**B**) The readily releasable pool (RRP) size was estimated from the charge released in the first 4 seconds of the response to 500 mM sucrose. Control n = 25/6, cDKO n = 26/6. (**C**) Vesicular release probability (Pves) was calculated as the ratio between the charge released by a single EPSC and the RRP. Control n = 24/6, cDKO n = 25/6. (**D**) The fraction of the RRP released by application of 250 mM sucrose. This is a measure for the fusogenicity of synaptic vesicles. Control n = 25/6, cDKO n = 26/6. (**E-F**) Results from fitting the sucrose traces to a minimal vesicle state model. Control n = 12 – 16/6, cDKO n = 7 – 18/6. See Figure S4 A for a scheme of the model. (**E**) The fitted maximal fusion rate constant (k_2,max_). 0 mM sucrose corresponds to mEPSC release. (**F**) The change in activation energy (ΔEa), reflecting the energy barrier for fusion. (**G-I**) The recovery of the first EPSC amplitude (1st) was tested by applying RRP-depleting trains (80 pulses at 40 Hz) and stimulating with a single pulse 2 seconds (R1) and 60 seconds (R2) after the train. Control n = 33/6, cDKO n = 38/6. (**G**) Example traces. (**H**) The amplitude of recovery pulse R1 was divided by the first amplitude of the train. (**I**) The absolute amplitude of recovery pulse R1. See Figure S4 J-K for quantifications of R2. (**J-M**) The recovery of the RRP was tested by dual applications of 500 mM sucrose within 30 s. Initial = first application. Recov = recovery application. cDKO n = 10/6, control n = 11/6. (**J**) Normalized example traces. (**K**) The percentage of the RRP replenished after 30 seconds. (**L**) The fusion rate and (**M**) the change in the activation energy of the initial and recovery applications. Note that the initial application corresponds to the 500 mM application shown in panels **A**-**F**. Control n = 12 - 16/6, cDKO n = 11 - 18/6. In **E**, **F**, **L** and **M**, data are presented as mean ± SEM. *p<.05, ***p<.001. Abbreviations: n.s. (not significant), HS (hypertonic sucrose), recov (recovery). See also Figure S5.

We have previously shown that hypertonic sucrose responses from autaptic neurons can be mathematically fitted using a minimal vesicle state model to calculate the fusion rate and the energy barrier for fusion ^72, 73^ (Figure S4A). To assess these parameters in tomosyn cDKO neurons, we applied this model to our sucrose responses. Model fitting revealed that neurons lacking tomosyns have strongly increased fusion rates at rest and during sucrose applications (Figure 2E). Accordingly, the energy barrier was strongly reduced in cDKO neurons compared to controls (Figure 2F). The fitting also yields estimates for the RRP size as well as the priming and fusion rates of SVs. These fitted estimates confirmed the unchanged RRP size and increased P_ves_ seen before (Figures S4B-D). Priming and de-priming rates were not significantly altered in the absence of tomosyns, although a trend to lower values was observed (Figures S4E and F). Overall, these results show that tomosyns do not influence the number of releasable vesicles, but instead increase the energy barrier for their fusion.

### Loss of tomosyns impairs synaptic recovery

To test whether tomosyns impact the recovery of synaptic strength after intense activity, single recovery pulses were given 2 seconds (R1) and 60 seconds (R2) after exhaustive high-frequency stimulation (Figure 2G). This also allowed for an alternative measure of the RRP by back-extrapolating the cumulative charge released by the stimulus train to the y-intercept (Figure S4G) ^74^. Again, the RRP size was similar in control and cDKO neurons (Figure S4H). In addition, the refilling rate of SVs was unchanged, measured by the slope of the back-extrapolation line (Figure S4I). Nevertheless, while control neurons recovered to ∼100% of their initial EPSC amplitude within 2 seconds, cDKO neurons only reached ∼60% (Figure 2H). Notably, the absolute EPSC amplitude was similar in control and cDKO neurons at this point (Figure 2I). During the second recovery pulse given after a full minute, cDKO neurons had restored their initially larger EPSC size (Figures S4J and S4K). To further explore this recovery phenotype, we performed a dual sucrose experiment to measure the extent of calcium-independent RRP recovery (Figure 2J). Both control and cDKO neurons recovered their RRP to 80% within 30 seconds (Figure 2K). Interestingly, the fusion rate and energy barrier after 30s recovery were similar to control levels, in stark contrast to the initial values (Figure 2L and M). Together, these results show that while SV refilling is normal, cDKO neurons take longer to recover their enhanced synaptic strength. This suggests that tomosyns reduce the initial release probability to maintain synaptic strength and accelerate synaptic recovery.

### Ultrastructural changes do not explain increased synaptic strength in tomosyn cDKO synapses

Nematode and fly tomosyn mutants were shown to have more docked vesicles, suggesting that invertebrate tomosyn inhibits SV docking ^39, 41^. To test whether tomosyns affect SV docking in mammalian neurons, we performed high-pressure freeze electron microscopy on low density cultures of cDKO neurons grown on glia-covered sapphires (Figures S5A). We observed no significant difference in the number of SVs contacting the active zone between control and cDKO synapses, except for a trend towards reduced SV docking in cDKO synapses (Figures S5A-B). Expressing tomosyn-1m, the most abundant tomosyn isoform in the hippocampus ^75^, did not affect SV docking in cDKO cultures (+WT), but the number of docked SVs was reduced compared to control neurons (Figures S5A-B). The total number of vesicles per synaptic profile, active zone length and vesicle distribution was normal (Figure S5C-E). These data show that the increased synaptic strength in tomosyn cDKO neurons is unlikely to be explained by an increased docked/primed vesicle pool in mammalian neurons.

### Tomosyn-syb2/VAMP2 hybrids fail to mimic tomosyn’s inhibitory function

The C-terminal SNARE motif of tomosyns is thought to compete with the corresponding motif in syb2/VAMP2 during SNARE assembly (Figure 3A). In such a scenario, tomosyns would be equally functional with the syb2/VAMP2 SNARE motif. To test this, a hybrid tomosyn was cloned in which the C-terminal SNARE motif of tomosyn-1m was replaced with the SNARE motif of syb2/VAMP2 (Figure 3B). Expressing wildtype tomosyn in cDKO neurons restored normal synaptic tomosyn levels, while hybrid tomosyn was expressed at an even higher level (Figures 3C and S6). Neuronal morphology was not altered by the expression of either variant and both variants targeted normally to synapses (Figure S6D). Expression of wild-type tomosyn-1m restored the enhanced synaptic depression in cDKO neurons (Figures 3D-F) as well as the defect in synaptic recovery after high-frequency stimulation (Figures 3G and 3H). Interestingly, hybrid tomosyn rescued these synaptic changes only to a limited extent (Figures 3D-H), despite the high expression levels of this variant. These observations are inconsistent with the simple competition model, in which tomosyns regulate synaptic transmission by competing with syb2/VAMP2 in SNARE assembly.

**Figure 3.**
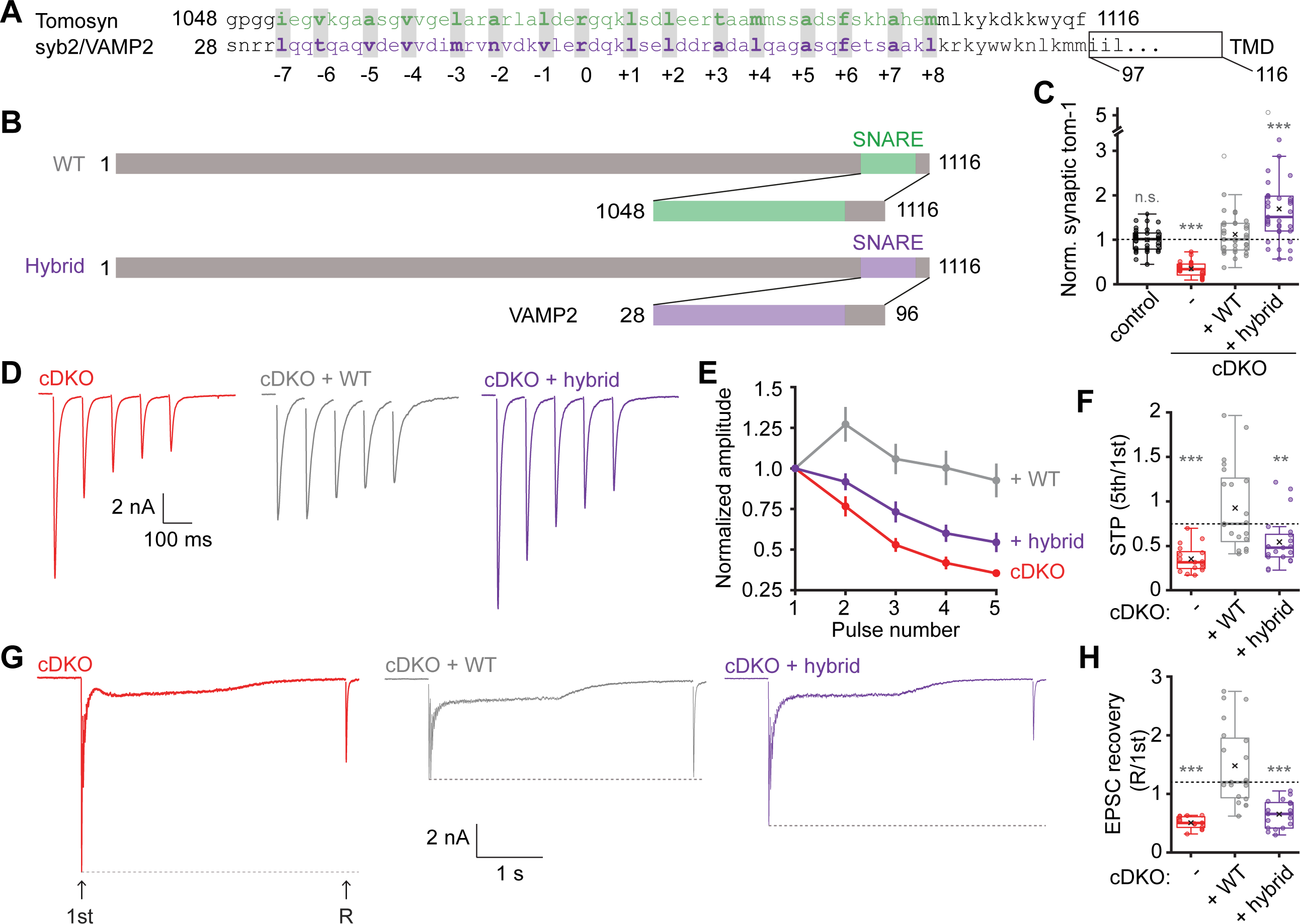
A tomosyn-VAMP2 hybrid fails to fully restore tomosyns inhibitory function. (**A**) Comparison of the amino acid sequences of the syb2/VAMP2 and tomosyn R-SNARE motifs. TMD = transmembrane domain. (**B**) Schematic representation of WT tomosyn-1m and the hybrid mutant, in which we replaced the C-terminus of tomosyn with the corresponding region in VAMP2. Numbers correspond to amino acids. (**C**) Synaptic tomosyn-1 expression from immunostainings, normalized to control within each neuronal culture. Control n = 35/3, cDKO n = 30/3, + WT n = 33/3, + Hybrid n = 35/3. See Figure S6 for example images and morphological analysis. (**D-F**) Analysis of short-term plasticity (STP) by stimulating neurons with 5 pulses at 10 Hz. cDKO n = 17/4, + WT n = 21/4, + Hybrid n = 21/4. (**D**) Example traces. (**E**) Amplitudes were normalized to the first pulse of the train. (**F**) STP quantified by the ratio of the fifth pulse over the first pulse. (**G-H**) Recovery of the first EPSC amplitude (1st) was tested by high-frequency stimulation (80 pulses at 40 Hz) followed by a recovery pulse after 2 seconds (R). cDKO n = 14/4, + WT n = 20/4, + Hybrid n = 18/4. (**G**) Example traces. (**H**) The amplitude of the recovery pulse was divided by the first amplitude of the train. In **E**, data are presented as mean ± SEM. Bonferroni-corrected alpha levels: panel **C**: ***p<.00033; panels **F** and **H**: **p<.005, ***p<.0005. Grey asterisks show comparison to + WT group. Abbreviations: TMD (transmembrane domain), n.s. (not significant).

### The tomosyn SNARE motif forms a stable ternary complex with syntaxin-1 and SNAP-25

We then set out to dissect how tomosyns may affect SNARE assembly. Previous experiments demonstrated that the SNARE motif of tomosyns forms a stable ternary complex with syntaxin-1 and SNAP-25 ^32^, but the energetics and kinetics of the complex formation have not been well measured. We previously developed a single-molecule assay based on high-resolution optical tweezers to characterize SNARE assembly ^8, 19, 76–78^. Here, we adopted the assay to measure the folding energy and kinetics of the tomosyn SNARE complex. As in our previous assay for the SNARE complex containing syb2/VAMP2, a single tomosyn SNARE complex was now connected to two polystyrene beads either directly or via a DNA handle at the C-termini of tomosyn and syntaxin-1 (Figure 4A). The two proteins were crosslinked at their −6 layers through a disulfide bond (the tomosyn-Syx conjugated at X-6) to facilitate reversible SNARE assembly and disassembly ^79^. The beads were held in two optical traps as force and displacement sensors. When the tomosyn SNARE complex was being pulled by moving one trap relative to the other, the extension and tension of the protein-DNA tether were measured to report conformational changes of the complex and their associated energies.

**Figure 4.**
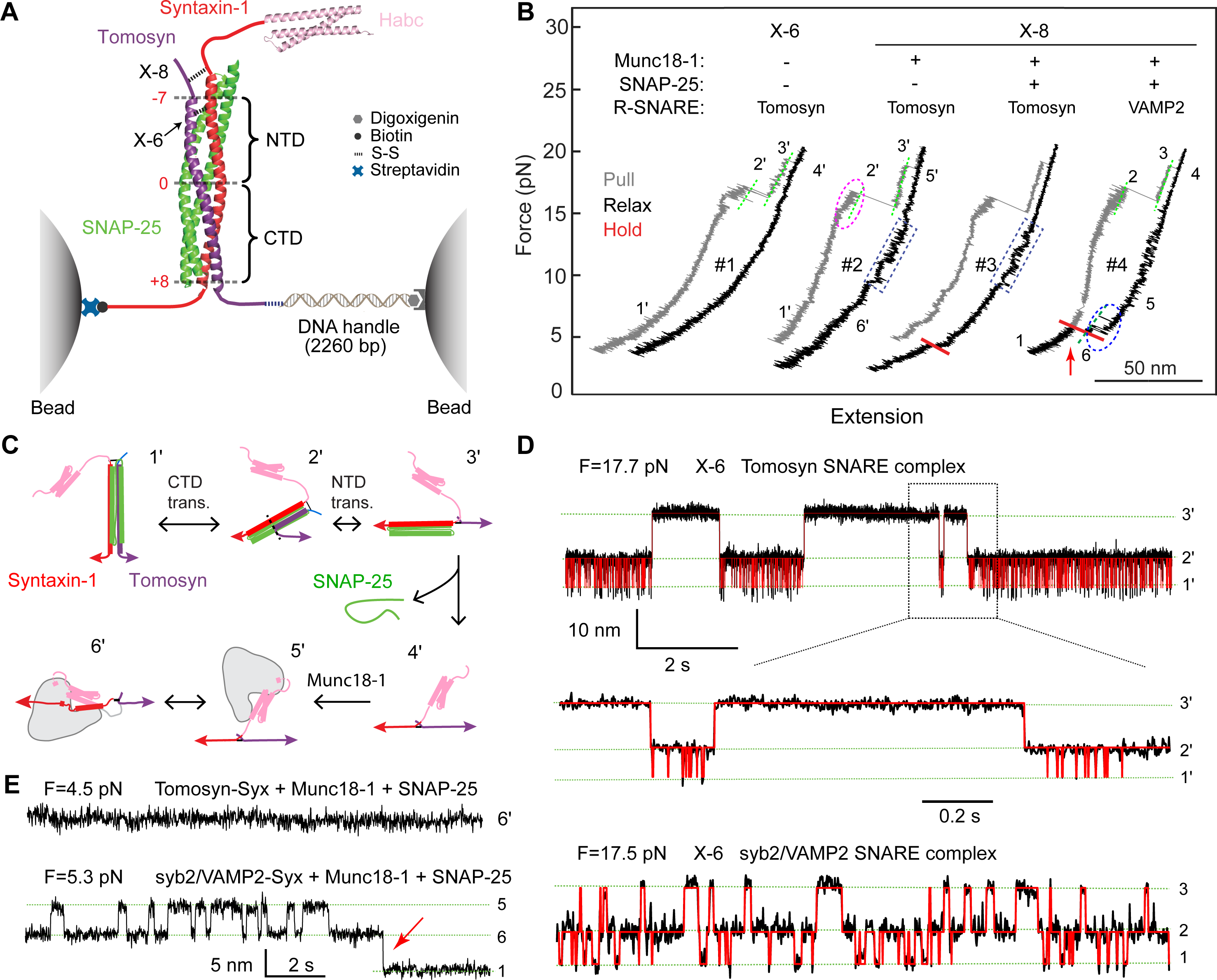
The SNARE motif of tomosyn forms a stable SNARE complex with syntaxin-1 and SNAP-25, but fails to form a template complex with Munc18-1 and syntaxin-1. (**A**) Experimental setup to pull a single tomosyn SNARE complex (PDB ID: 1URQ) using optical tweezers. The tomosyn and syntaxin-1 (Syx) molecules in the complex were attached to two polystyrene beads via a DNA handle at their C-termini and crosslinked at either the −6 layer (X-6) or the −8 layer (X-8) as indicated. The central ion layer divides the parallel helix bundle into the N-terminal domain (NTD) and the C-terminal domain (CTD). Munc18-1 and SNAP-25 may be added to the solution either alone or together to test template complex formation or SNARE assembly. (**B**) Force-extension curves (FECs) obtained by pulling (gray) or relaxing (black) the tomosyn-Syx conjugate at a speed of 10 nm/s or holding it at a constant mean force or trap separation (red). Different FEC regions, which may be indicated by green dashed lines, are labeled by the numbers of their associated states in C. (**C**) Schematic diagrams of different states 1’-6’ involved in SNARE folding/assembly and Munc18-1 association. See also Figure 5C for states 1-7. (**D**) Extension-time trajectories at the indicated constant forces (F) showing three-state folding and unfolding transitions of the tomosyn or syb2/VAMP2 SNARE complex. Red curves represent the idealized transitions derived from hidden Markov modeling. Green dashed lines mark the average extensions of the associated states labeled on the right. (**E**) Extension-time trajectories of different SNARE conjugates at constant low forces.

When pulled to a high force, the tomosyn SNARE complex unfolded stepwise with characteristic intermediate states and kinetics as seen in the force-extension curve (FEC, Figure 4B, FEC#1, grey curve). The FEC is quantitatively similar to that of the syb2/Vamp2 SNARE complex, which helps identify the intermediate states ^79^. These states include a half-zippered SNARE bundle (Figure 4C, state 2’) and the unzipped tomosyn SNARE motif and the binary t-SNARE complex syntaxin-1:SNAP-25 (state 3’) ^80^, and completely unfolded tomosyn-Syx conjugate and dissociated SNAP-25 (state 4’). Relaxing the unfolded tomosyn-Syx conjugate did not cause refolding of the remaining SNAREs. Thus, the tomosyn SNARE motif barely associates with syntaxin-1 in the absence of SNAP-25. This observation also suggests that, like the syb2/VAMP2 SNARE complex, the tomosyn SNARE complex requires the t-SNARE complex as an intermediate for its assembly ^8, 80^.

We then held the tomosyn SNARE complex at constant trap separations or mean forces and detected its folding and unfolding transitions with high spatiotemporal resolution (Figure 4D). These trajectories revealed reversible transitions among the folded four-helix bundle state (state 1’), the half-zippered state 2’, and the unzipped state 3’. Hidden-Markov analyses of the trajectories demonstrated that the transitions are sequential and direct transitions between states 1’ and 3’ are negligible ^81^. Thus, the folding and unfolding transitions of the tomosyn SNARE complex can be divided into the transitions of the C-terminal domain (CTD) and the N-terminal domain (NTD) (Figure 4A). The CTD transition is frequent, indicating a great transition rate, while the NTD transition is ∼400-fold slower (Figure 4D). Based on the measured force ranges for both transitions, we estimated unfolding energies of 23 (±1, SEM) k_B_T and 35 (±2) k_B_T for the CTD and the NTD, respectively, compared with the corresponding CTD energy of 22 k_B_T and NTD energy of 38 k_B_T for the syb2/Vamp2 SNARE complex ^79^. Overall, the tomosyn SNARE complex folds and unfolds similarly to the syb2/Vamp2 SNARE complex (Figure 4D), including their similar energetics ^76, 79^. However, the NTD of the tomosyn SNARE complex folds much slower than that of the syb2/Vamp2 SNARE complex.

### The tomosyn SNARE motif does not form a template complex with Munc18-1 and syntaxin-1

We and others have shown that for physiological SNARE assembly, syntaxin-1 and syb2/VAMP2 first bind on the surface of Munc18-1 to form a ternary template complex, which then binds SNAP-25 to conclude SNARE assembly and displace Munc18-1 from the folded four-helix bundle ^8, 17, 19, 23^. To examine whether tomosyn can replace syb2/VAMP2 in the template complex, we crosslinked tomosyn and syntaxin-1 at their −8 layers (X-8) as in our previous assay ^19^. Crosslinking at this alternative site opens the closed syntaxin-1 conformation bound by Munc18-1 and minimally perturbs the template complex. When the unfolded tomosyn-Syx conjugate was being slowly relaxed in the presence of Munc18-1 in the solution, Munc18-1 first bound to syntaxin-1, forming open syntaxin-1 in a reversible manner in the force range of 10-17 pN as previously observed (Figure 4B, FEC#2 & #3 in blue dashed rectangles; Figure 4C, state 6’). However, no further folding transition was detected at a lower force range, as is expected for the formation of a template complex (Figure 4B, #2, black FEC), even when the conjugate was held at a low force for a long time (Figure 4B, #2, red region). Consequently, no SNAP-25 binding and SNARE assembly was observed (N=23, Figure 4B, #3). For comparison, the syb2/VAMP2-Syx conjugate forms the template complex in the force range of 3-7 pN with an extension change of 5.4 nm (Figure 4B, #4, blue dashed oval), which frequently bound to SNAP-25 in the solution to form the ternary SNARE complex (the extension-drop indicated by the red arrow in Figure 4b, #4). These comparisons were confirmed by the experiments at a constant mean force (Figure 4E). In conclusion, the tomosyn SNARE motif failed to form a template complex with Munc18-1-bound open syntaxin-1 to chaperone SNARE assembly.

### The tomosyn SNARE motif induces a large conformational change in the template complex

To further explore the molecular mechanism underlying tomosyn’s function, we examined whether the tomosyn SNARE motif modulates the formation of the synaptic template complex using the syb2/VAMP2-Syx conjugate (Figure 5A). We repeated the above template complex assay in the presence of 2 µM Munc18-1 and 2 µM tomosyn SNARE motif in the solution (Figures 5B and 5C). Surprisingly, an intermediate state 7* frequently appeared in the otherwise binary transition between the template complex state 7 and open syntaxin-1 state 6 (compare the first and second trajectories in Figure 5D), with an average extension ∼1.5 nm greater than that of the template complex state and ∼4 nm less than that of open syntaxin-1 (Figure S7). This new state is syb2/VAMP2-dependent, as the tomosyn SNARE motif did not induce further folding of Munc18-1-bound open syntaxin-1 when either the tomosyn-syntaxin-1 conjugate (Figures 4B & 4E) or syntaxin-1 alone (Figures 5E & S8) was being pulled. These observations suggest that the tomosyn SNARE motif specifically binds to the template complex to induce a large conformational change in the template complex as represented by its 1.5 nm extension change.

**Figure 5.**
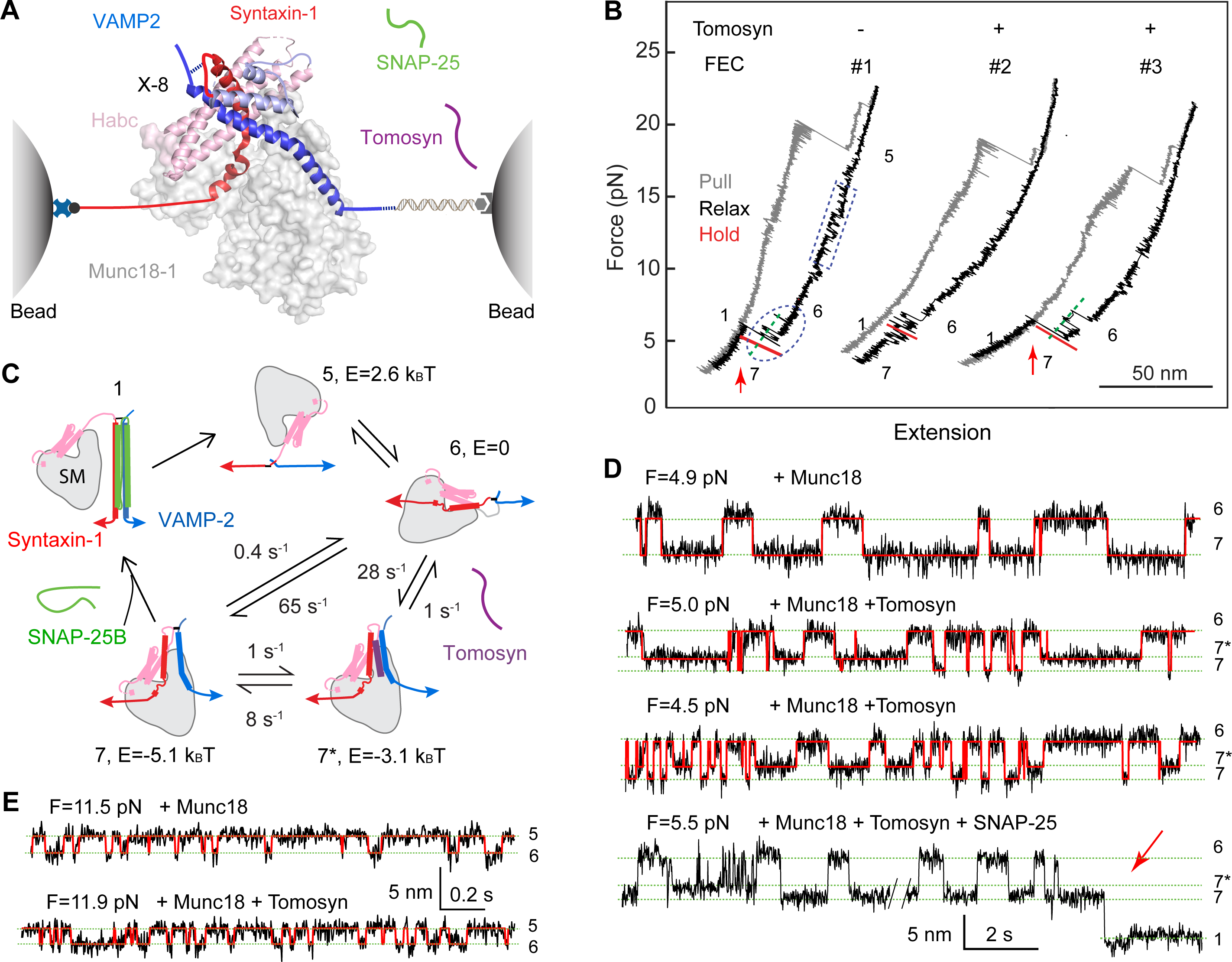
The tomosyn SNARE motif binds to the template complex to block its association with SNAP-25. (**A**) Experimental setup to investigate the effect of the tomosyn SNARE motif on the folding of the synaptic template complex Munc18-1:syntaxin-1:VAMP2 (PDB ID 1SFC). (**B**) FECs of the VAMP2-Syx conjugate in the presence of 2 µM Munc18-1, 60 nM SNAP-25, and 0 (#1) or 2 µM (#2 and #3) tomosyn SNARE motif in the solution. The dashed rectangle and oval mark reversible folding and unfolding transitions of Munc18-1-bound open syntaxin and the template complex, respectively. Red arrows indicate events of SNAP-25 binding to the template complex. (**C**) Schematic diagrams of different states and their associated energies (E) and transition rates. (**D**) Extension-time trajectories at the indicated constant forces (F) showing folding and unfolding transitions between the states labeled on the right. The red curves represent the idealized transitions derived from hidden Markov modeling. The red arrow designates the SNAP-25 binding event. (**E**) Extension-time trajectories at constant forces showing folding and unfolding transitions of the Munc18-1-bound open syntaxin-1 in the absence (top) or presence of the tomosyn SNARE motif in the solution.

We characterized the stability and folding kinetics of the tomosyn-bound template complex. To this end, we extensively measured the three-state transitions at different constant mean forces and analyzed the resulting time-dependent extension trajectories using hidden Markov modeling ^81^ (Figure 5D). The analyses revealed the probabilities and lifetimes of the three states and their associated transition rates. Extrapolation of these force-dependent quantities to zero force using an energy landscape model of protein folding yielded the unfolding energies of the tomosyn-bound and unbound template complexes to be 3.1 (± 0.1, S.E.) k_B_T and 5.1 (± 0.2) k_B_T, respectively (Figure 5C) ^82^. The unfolding energy of the template complex is consistent with our previous measurement of 5.2 (± 0.2) k_B_T ^19^. The average lifetimes of these states are ∼ 0.7 s and ∼ 1.6 s for the tomosyn-bound and unbound template complexes, respectively, as well as ∼ 0.014 s for the Munc18-1-bound open syntaxin-1. In the presence of 2 µM tomosyn SNARE motif in the solution, the tomosyn-bound template complex efficiently forms from two pathways, either coupled binding of tomosyn and syb2/VAMP2 to Munc18-1-bound open syntaxin-1 with a rate of ∼ 28 s^−1^, or direct binding of tomosyn to the pre-formed template with a rate of ∼ 1 s^−1^. In addition, the tomosyn SNARE motif dissociates from the template complex with a rate of 8 s^−1^. Compared to the direct folding rate of 65 s^−1^ for the template complex, the tomosyn-bound template complex is expected to compete with the template complex for its formation, thereby modulating Munc18-1-chaperoned SNARE assembly.

### The tomosyn-bound template complex does not bind SNAP-25

To uncover the function of the tomosyn-bound template complex in SNARE assembly, we tested its binding to SNAP-25 during the three-state transitions in the presence of 60 nM SNAP-25 in the solution. Interestingly, SNAP-25 only binds to the template complex (Figure 5C, state 7) ^19^, but not the tomosyn-bound template complex (state 7*), to form the SNARE four-helix bundle (N=28, Figure 5B, #3; Figure 5D, bottom trace). Due to the presence of the tomosyn-bound template complex, the probability to observe chaperoned SNARE assembly within ∼100 s detection time was reduced to 0.13, compared to the probability of 0.7 in the absence of tomosyn ^19^. Thus, tomosyn attenuates chaperoned SNARE assembly by binding to the template complex to inhibit its association with SNAP-25.

### A polybasic domain outside the SNARE motif is critical for tomosyn function

The above findings prompted us to re-evaluate the importance of the different functional domains in tomosyn, starting with a point mutation in the +6 layer of the SNARE motif (FA-mutant, Figure 6A). This mutant was based on the F77A mutant in syb2/VAMP2 which abolished template complex formation, chaperoned SNARE assembly and secretory vesicle fusion ^79, 83, 84^. In contrast to wild-type tomosyn, the FA-mutant failed to reverse the enhanced synaptic depression in cDKO neurons (Figures 6B and 6C) or restore synaptic recovery after high-frequency stimulation (Figures 6D and 6E). However, while the FA-mutant was targeted to synapses, its expression level was lower than wild-type tomosyn, which could partly contribute to its reduced rescue ability (Figures 6F, 6G and S9).

**Figure 6.**
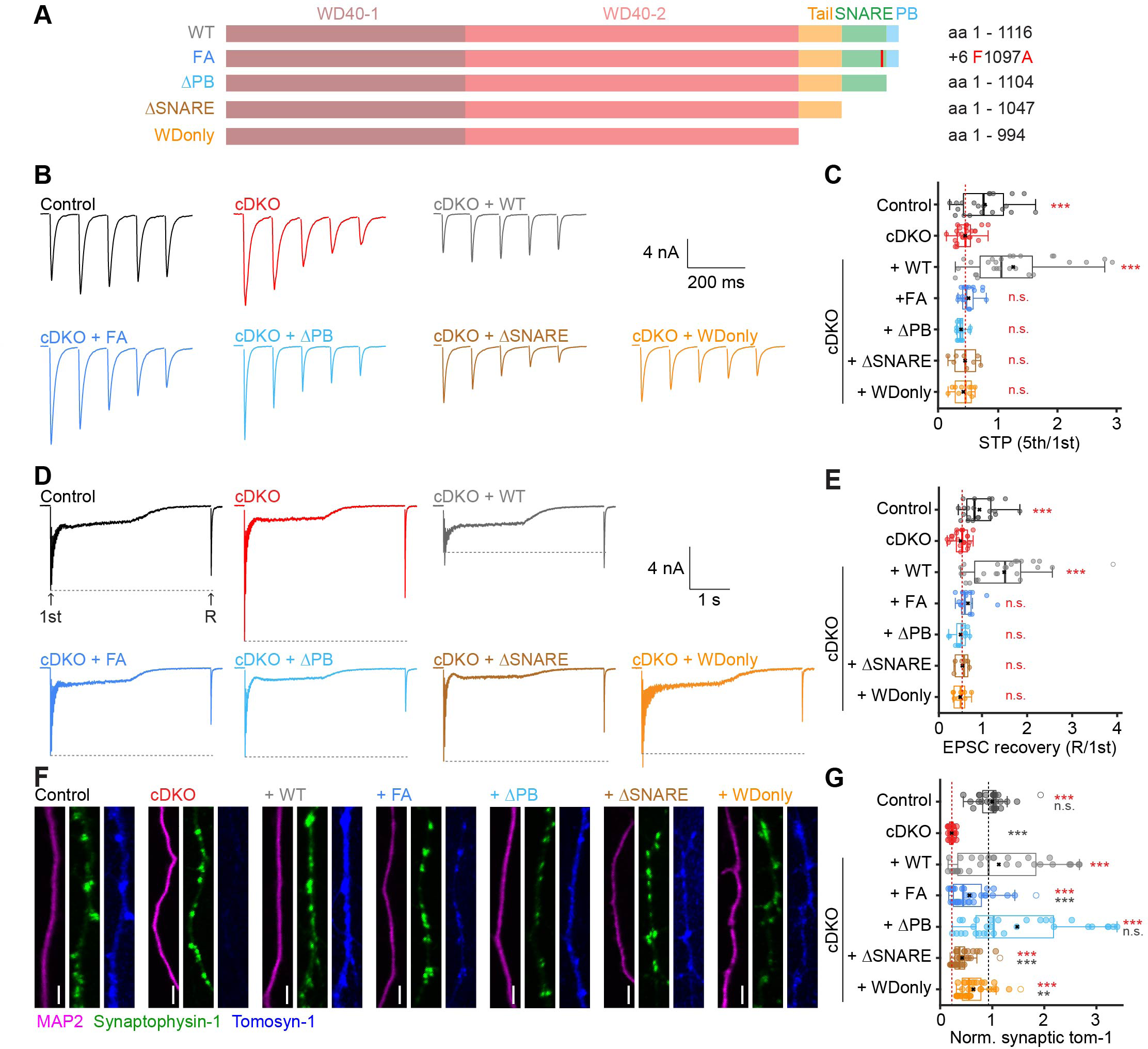
A polybasic domain outside the SNARE motif is critical for tomosyns function. (**A**) Schematic representations of WT tomosyn-1m and mutants created to test its individual domains. The amino acid (aa) numbers used for each construct or its mutation site are stated at the right. (**B-C**) Short-term plasticity (STP) was measured during 5 pulses at 10 Hz. Control n = 21/6, cDKO n = 25/8, +WT n = 27/6, +FA n = 18/5, +ΔPB n = 13/4, +ΔSNARE n = 8/4, +WDonly n = 10/3. (**B**) Example traces. (**C**) STP quantified by the ratio of the fifth pulse over the first pulse of the train shown in **B**. (**D-E**) Recovery of the first EPSC amplitude (1st) was tested by high-frequency stimulation (80 pulses at 40 Hz) followed by a recovery pulse after 2 seconds (R). Control n = 21/6, cDKO n = 25/8, +WT n = 25/6, +FA n = 16/5, +ΔPB n = 12/4, +ΔSNARE n = 6/3, +WDonly n = 9/2. (**D**) Example traces. (**E**) The amplitude of the recovery pulse was divided by the first amplitude of the train. (**F-G**) Immunostainings to confirm expression of all mutants. Control n = 26/3, cDKO n = 27/3, +WT n = 29/3, +FA n = 33/3, +ΔPB n = 30/3, +ΔSNARE n = 36/3, +WDonly n = 34/3. (**F**) Example images of dendrites. Scale bar = 10 µm. (**G**) Synaptic tomosyn-1 expression normalized to control within each neuronal culture. See Figure S9 for whole-neuron images and morphological analysis. Bonferroni-corrected alpha levels: **p<.00166, ***p<.000166. Red asterisks show comparison to cDKO, grey asterisks show comparison to + WT group. Abbreviations: PB (polybasic domain), n.s. (not significant).

Tomosyns contain two N-terminal WD40 double propeller regions (together referred to as WD40 domains), a proposed auto-inhibitory tail domain ^85–87^, a SNARE motif and a polybasic domain at the COOH-terminal end that contains multiple lysines (Figure 6A). We next tested a series of C-terminal truncations. The first truncation has an intact SNARE motif, but lacks the adjacent polybasic domain (ΔPB, Figure 6A). Surprisingly, the ΔPB-mutant failed to restore the cDKO phenotype (Figures 6B-E), even though this mutant was expressed at similar levels as wild-type tomosyn (Figures 6F, 6G and S9). Two consecutive larger truncations either removed the SNARE motif (ΔSNARE-mutant; Figure 6A), or the SNARE and tail domain, leaving only the WD40 domain intact (WDonly-mutant; Figure 6A). Again, expression of both these mutants could not restore the cDKO phenotype (Figures 6B-E), suggesting that the WD40 domains alone are not sufficient for tomosyn function, irrespective of the presence of the tail domain. Notably, just as the FA-mutant, these mutants were properly targeted to synapses but their expression levels were reduced compared to wild-type tomosyn (Figures 6F, 6G and S9). Hence, synaptic tomosyn protein levels might depend on the presence of a functional SNARE motif. Taken together, these results suggest that tomosyns require both an intact SNARE motif and the adjacent polybasic domain to function properly.

### The polybasic domain contributes to the reduced functionality of the syb2/VAMP2-hybrid

The C-terminal region targeted in the tomosyn-syb2/VAMP2 hybrid tested above spanned both the SNARE motif and polybasic domain (Figure 3). To test whether one domain was solely responsible for the reduced functionality, partial hybrid constructs were designed: a core-hybrid in which only the SNARE motif was replaced and a linker region (LR)-hybrid in which the linker region of syb2/VAMP2 replaced the polybasic domain of tomosyn (Figure 7A). These tomosyn variants were properly targeted to synapses, were expressed at similar levels and did not alter neuronal morphology (Figures 7B and S10). While wild-type tomosyn efficiently restored the defects in short-term plasticity and synaptic recovery in cDKO neurons, both the core-hybrid and LR-hybrid only partially rescued these synaptic defects (Figures 7C-G), similar to our previous findings. Thus, the core SNARE motif and the adjacent polybasic domain both contributed to the reduced functionality of the original hybrid mutant, confirming the requirement of both domains for tomosyn function.

**Figure 7.**
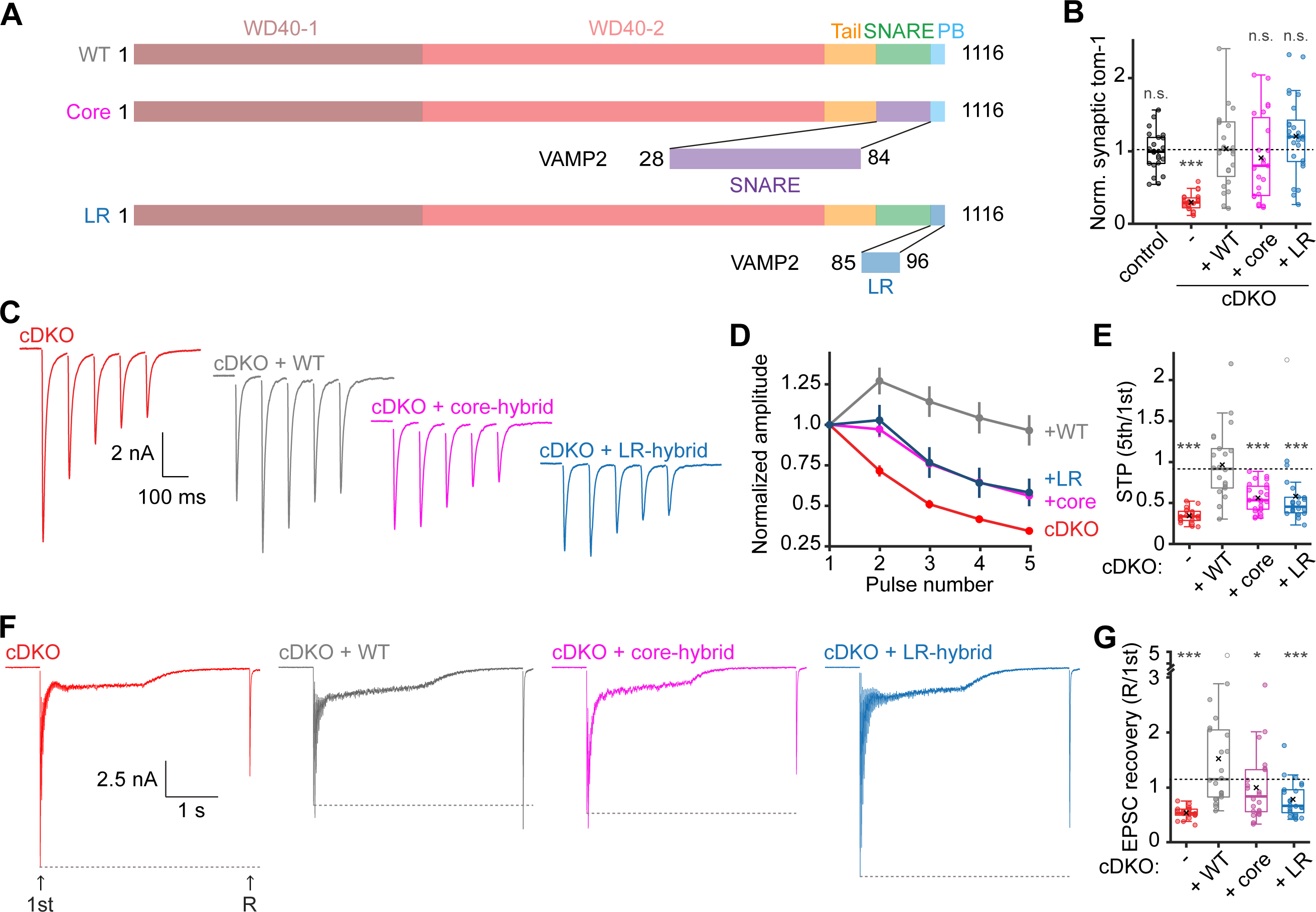
The polybasic domain contributes to the reduced functionality of the VAMP2-hybrid. (**A**) Schematic representation of mutant constructs created to test the individual contributions of the linker region and the SNARE motif to the lack of interchangeability of the corresponding regions in tomosyn and VAMP2. (**B**) Normalized synaptic tomosyn-1 expression. Control n = 22/4, cDKO n = 23/4, + WT n = 24/4, + Core n = 23/4, + LR n = 25/4. See Figure S108 for example images and morphological analysis. **C-E**) Short-term plasticity (STP) was estimated by stimulation with 5 pulses at 10 Hz. cDKO n = 20/4, + WT n = 22/4, + Core n = 24/4, + LR n = 23/4. (**C**) Example traces. (**D**) Amplitudes were normalized to the first pulse. (**E**) STP is quantified by the ratio of the fifth over the first amplitude. (**F-G**) Recovery of the initial amplitude was tested by high-frequency stimulation (80 pulses at 40 Hz) followed by a recovery pulse after 2 seconds. cDKO n = 19/4, + WT n = 22/4, + Core n = 22/4, + LR n = 22/4. (**F**) Example traces. (**G**) The amplitude of the recovery pulse was divided by the first amplitude of the train. In **D**, data are presented as mean ± SEM. Bonferroni-corrected alpha levels: *p<.0166, ***p<.00033. Grey asterisks show comparison to + WT group. Abbreviations: PB (polybasic domain); LR (Linker region); n.s. (not significant).

## Discussion

In this study, we characterized the role of the two mammalian tomosyn genes in SNARE complex assembly, SNARE-dependent membrane fusion and synaptic transmission. Hippocampal neurons lacking both tomosyns exhibited a decreased energy barrier for synaptic vesicle fusion and, consequently, an increased initial synaptic strength, followed by more pronounced synaptic depression and slower recovery (Figures 1 and 2). While expression of wild-type tomosyn-1 fully rescued cDKO phenotypes, a hybrid containing the SNARE motif of syb2/VAMP2 did not (Fig. 3). In line with these findings, single molecule force measurements revealed that tomosyn’s SNARE motif fails to form a template complex with Munc18-1 and syntaxin-1 (Figure 4), but instead binds to the synaptic template complex VAMP2:Munc18-1:Syntaxin-1 to block its association with SNAP-25 (Figure 5). Finally, rescue experiments revealed that tomosyns’ complete C-terminus is required for its effects on synaptic transmission (Figures 6 and 7).

Our data show that tomosyn deficiency leads to a substantially increased initial synaptic strength. This is consistent with previous findings in nematode and fly neuromuscular junction ^39–41^. However, in these studies, tomosyn deficient synapses had larger docked and primed vesicle pools. In contrast, we found no evidence for enlargement of docked/primed pools. Instead, our results indicate that mammalian tomosyns control membrane fusion by increasing the energy barrier for fusion (Figures 2), leading to decreased probability that vesicles fuse upon stimulation (P_ves_). This is consistent with a previous study in the mouse neuromuscular junction, where synapses lacking tomosyn-2 had a high spontaneous release frequency and exhibited faster synaptic depression ^47^. Synaptic vesicles are known to have heterogeneous P_ves_ (e.g., primed and superprimed vesicles) ^16, 88–92^. Tomosyns may shift the balance towards more low-P_ves_ vesicles, causing the same stimulus to release a smaller fraction of the vesicles and thereby limiting initial synaptic strength, but also limiting synaptic depression (Figures 8). Because tomosyns do not influence the vesicle recruitment/priming rate (Figure 2K and S4I), their net effect is to allow vesicle fusion to be more evenly distributed during action potential trains. These short-term plasticity phenomena are regarded as crucial in synaptic computation, such as working memory ^93^ and sensory processing ^25^.

The effect of tomosyn deficiency on synaptic strength resembles the effects of phorbol ester application ^68, 72^, Munc13 activation ^71, 94, 95^, PKC-dependent phosphorylation of Munc18-1 ^8, 96^, Synaptotagmin-1 ^97^, or Liprin-alpha ^98^, or synaptic plasticity phenomena such as augmentation ^69^ and post tetanic potentiation (PTP) ^73^. These effects/phenomena are all explained by higher P_ves_. Tomosyns act in the opposite direction, reducing P_ves_.

While tomosyns reduce synaptic strength at the onset of activity, synaptic responses following exhaustive stimulation were similar in control and tomosyn-deficient neurons (Figure 2G-I). Under these conditions, most fusing vesicles are probably newly recruited vesicles. These vesicles may not yet be targeted by tomosyns. This is further supported by the observation that tomosyns only reduced the energy barrier for vesicles released during the initial sucrose application, but not in newly recruited vesicles that were released during a second application (Figure 2L-M). Many studies indicate that new vesicles have a low release probability and take seconds to gain a higher release probability (‘super-primed’) ^91, 99–101^, possibly as a consequence of the time required to assemble mature release machinery. Tomosyns may primarily inhibit the transition to this mature, high-P_ves_ state(s).

Several molecular explanations for different P_ves_ among vesicles have been postulated (discussed in ^16^), including (i) their different SNARE assembly state ^14, 102^, (ii) a variable coupling between vesicles and Ca^2+^-channels ^103–105^, and/or (iii) the variable number of SNARE complexes per vesicle ^106^. Our data are consistent with each of these scenarios: (i) The force measurements are consistent with a two-step priming scheme where tomosyns reduce the fraction of synaptic vesicles equipped with mature release machinery ^13^; (ii) Tomosyns were detected among the core interactome of Ca^2+^_-_ channels ^107^. However, tomosyns also influence P_ves_ upon Ca^2+^-independent stimuli (sucrose application, Figure 2), indicating that Ca^2+^-channel coupling cannot be the only explanation; (iii) By binding to at least a subset of assembling SNARE complexes per vesicle, tomosyns may create a situation where fewer SNAP-25-containing SNARE complexes per vesicle exert force and hereby create lower P_ves_ vesicles (Figure 8).

**Figure 8.**
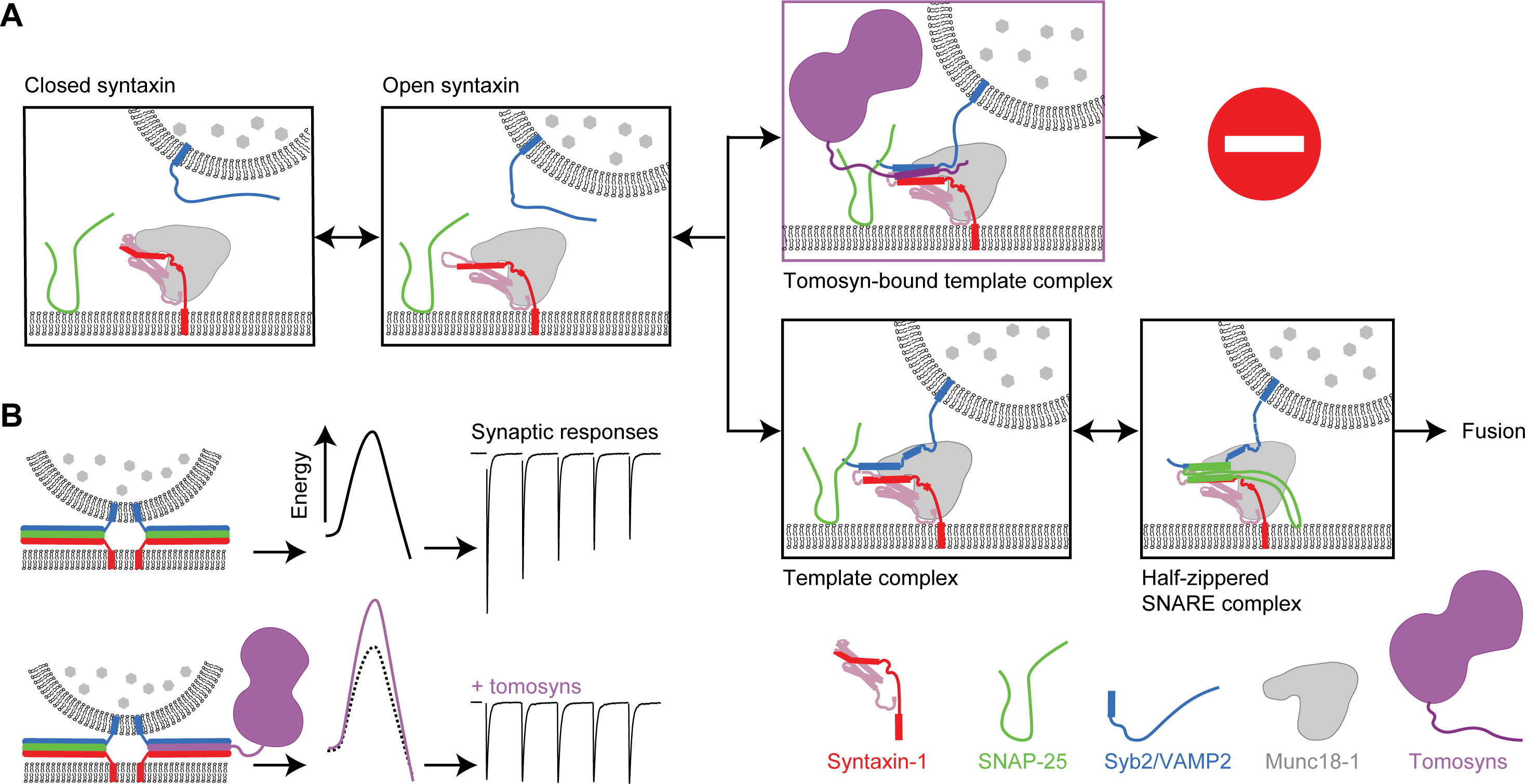
Graphic representation of the working model of tomosyn attenuating SNARE assembly and synaptic depression. (**A**) Munc18-1 sequesters syntaxin in a ‘closed’ conformation unable to associate with other SNAREs ^112^. This closed dimer transits to an open conformation in which the SNARE motif is accessible, assisted by Munc13 ^113^. Syb2/VAMP2 binds to the open-syntaxin complex to form the template complex in which the N-terminal regions of both SNARE motifs are aligned. Finally, tomosyns bind the VAMP2-containing template complex, preventing SNAP-25 binding and further SNARE assembly. (**B**) Multiple trans-SNARE complexes cooperate to overcome the fusion barrier in a manner dependent on their copy number and zippering states. Tomosyns bind the assembling template complex to produce a higher barrier SNARE complex state lacking SNAP-25. As a result, a smaller fraction of the vesicle pool will be released upon stimulation, which reduces synaptic strength and limits synaptic depression.

Our data argue against the concept that tomosyns directly compete with syb2/VAMP2 in SNARE assembly. This view is mainly derived from the observation that the tomosyn SNARE motif forms a stable ternary complex with syntaxin-1 and SNAP-25, similar to syb2/VAMP2 ^32^. We showed that, despite its much slower folding rate, the SDS-sensitive tomosyn SNARE complex is as thermodynamically stable as the SDS-resistant synaptic SNARE complex (Figure 4). Thus, our single molecule data generally support the previous observation on the stability of the tomosyn SNARE complex. However, two main findings challenge the *in vivo* relevance of this ternary complex: First, swapping the SNARE motif of tomosyn with that of syb2/VAMP2 failed to rescue tomosyn null mutant phenotypes, suggesting that the SNARE motif of tomosyns does not directly compete with syb2/VAMP2 for SNARE assembly (Figure 3). Second, tomosyn’s SNARE motif failed to form a template complex with Munc18-1 and syntaxin-1 in the way that syb2/VAMP2 does (Figure 5). Consequently, Munc18-1 did not promote the assembly of tomosyn-containing SNARE complexes, consistent with a previous observation ^31^. Taken together, our results challenge the proposed competition between tomosyns and VAMP2 during physiological SNARE complex assembly. Our data reveal that, instead, the SNARE motifs of both tomosyns and syb2/VAMP2 cooperatively bind to Munc18-1 bound open syntaxin-1 to form a tomosyn-bound template complex that blocks further SNARE assembly by association of SNAP-25.

This study provides further evidence that physiological SNARE assembly is mediated by the template complex ^17–19, 21–23, 108^. The template complex likely serves as a key target to regulate SNARE-mediated membrane fusion, for example via Munc13-1 ^109^ and phosphorylation of Munc18-1 ^19^. The template complex is stabilized by extensive interactions between Munc18-1 and SNAREs ^19, 23^, as well as interactions between the two aligned SNAREs. These interactions enhance the specificity of SNARE pairing. Thus, it is not surprising that tomosyns bind Munc18-1 in a way different from VAMP2, despite their similar SNARE motifs. In addition, both syb2/VAMP2 and syntaxin-1 are recruited by Munc13-1 and other regulatory proteins to form template complexes, and this recruitment depends on the membrane anchoring of both SNAREs ^6, 8, 109^. These observations may explain why tomosyn hybrids containing the SNARE motif of syb2/VAMP2 do not compete with vesicle-anchored VAMP2 for template complex formation (Figure 6). In conclusion, tomosyns target the VAMP2-containing template complex to regulate SNARE assembly and synaptic transmission.

Our results demonstrate that the inhibitory function of tomosyns requires the C-terminal polybasic domain (Figures 6 and 7). This domain may penetrate into the membrane-cytoplasm interface ^32^ and control the availability of the SNARE motif, as suggested for syb2/VAMP2 ^110^. Alternatively, the SNARE motif and the polybasic domain of tomosyns might cooperatively interfere with C-terminal zippering of partially assembled SNARE complexes into the juxtamembrane domains of syb2/ VAMP2 and syntaxin-1, which is proposed to deliver the power stroke in vesicle fusion ^76^. Reducing this zippering force, either by posing an additional barrier or by rendering a subset of SNARE complexes non-fusogenic, may reduce P_ves_ without affecting the readily releasable pool. The large N-terminal of tomosyns probably exerts a regulatory role in the inhibitory function of the SNARE motif^41, 57, 63, 111^.

In conclusion, our data show that tomosyns play a critical role in regulating synaptic strength, preventing primed synaptic vesicles from depletion during the onset of action potential trains (Figure 8). In this way, tomosyns limit synaptic depression and stabilize information transfer during repetitive firing. The exact impact of this regulation for cognitive functions remains to be determined, but modelling of working memory and the association of tomosyn mutations with neurodevelopmental disorders both suggest that this impact is substantial. More experiments are required to elucidate the structure of the tomosyn-bound template complex and the impact of tomosyn’s N-terminal domain on the structure.

## Supporting information

Supplemental Statistics Table item 1

## Acknowledgments

We would like to thank Joke Wortel for animal breeding, Ingrid Saarloos for cloning, Robbert Zalm for producing viral particles, Lisa Laan and Desiree Schut for preparing glia cultures and Joost Hoetjes for genotyping. We acknowledge Rien Dekker for high-pressure freeze electron microscopy. Furthermore, we would like to thank Vincent Huson for providing us with code and for assistance with fitting of hypertonic sucrose traces. We thank Niels Cornelisse, Ruud Toonen and Jacob Sørensen for their helpful discussion and comments on this work. This work is supported by the ZonMw-Veni program (09150161810052 to M.M.) from the Dutch Research Council (NWO), by the ERC Advanced Grant (322966) of the European Union to M.V. and NWO Gravitation program grant BRAINSCAPES (NWO 024.004.012) to M.V., the Horizon 2020 grant COSYN (RIA grant agreement no 610307, to M.V.), the Lundbeck Foundation Grant R277-2018-802 to M.V., the DFG (German Research Foundation) postdoctoral fellowship to A.S. (DFG project number SU 1131/1-1) and the NIH grant R35 GM131714 to Y.Z.

## Author contributions

M.M, M.Ö., J.Y., Y.Z. and M.V. conceptualized the study. M.M. and M.Ö. conducted and analyzed electrophysiology and immunofluorescence experiments. J.Y., A. K., and Y.Z. performed single molecule force measurements and analyzed the data, A.S. performed and analyzed Western blots. J.v.W. supervised and analyzed electron microscopy experiments. A.J.G. designed conditional tomosyn mice and performed initial characterizations. M.M., M.Ö., J. Y., Y.Z. and M.V. wrote the manuscript with input from all other authors.

## STAR Methods

### Animals

All animals were housed and bred according to institutional and Dutch Animal Ethical Committee regulations. The generation of homozygous Tom2^lox^ mice containing *LoxP* recombination sites flanking exon 3 of *Stxbp5l* was previously described ^47^. A similar strategy was used to generate Tom1^lox^ mice (Cyagen Biosciences). In short, a tomosyn-1 knock-in targeting vector (Tom1lox) was constructed by flanking exon 2 with *lox2272* recombination sites. Homologously targeted Tom1^lox^ embryonic stem cells (C57BL/6) were injected into blastocysts and implanted into pseudopregnant females to produce germ-line chimeras, which were mated with inbred C57BL/6 mice. Tom1/2 double lox mice were obtained by mating C57Bl/6 Tom1^lox^ mice with C57Bl/6 Tom2^lox^ mice. Newborn pups from homozygous matings were used for the preparation of neuronal cultures in all the described experiments.

### Cell culture

Animals were sacrificed on postnatal day 1 (P1). Hippocampi were dissected in cold Hanks’ balanced salt solution (Sigma, H9394) supplemented with 10mM HEPES (Gibco, 15630-056) and digested with 0.25% Trypsin (Gibco, 15090-046) for 10-20 min at 37°C. After washing three times with Hanks-HEPES the hippocampi were triturated in DMEM-glutamax (Gibco, 31966-021) supplemented with fetal calf serum (Gibco, 10270), non-essential amino acids (Sigma, M7145) and Penicillin-Streptomycin (Gibco, 15140-122) using fire-polished glass pipettes. For functional analyses, neurons were plated at a density of 2000 cells/well on 18 mm glass coverslips in 12-well plates. Coverslips contained micro-islands of freshly prepped rat glia in neurobasal medium (Gibco, 21103-049) supplemented with 2% B-27 (Gibco, 17504-044), 1.8% HEPES, 0.25% glutamax (Gibco, 35050-038) and 0.1% Penicillin-Streptomycin. Micro-islands were generated as described previously ^61^. Briefly, etched glass coverslips coated with agarose (Type II-A; Sigma, A9918) were stamped with 0.1 mg/mL poly-D-lysine (Sigma, P6407) mixed with 0.7 mg/mL rat tail collagen (BD Biosciences, 354236) and 10 mM acetic acid (Sigma, 45731) using a custom-made stamp. Freshly cultured rat glia were plated at a density of 8000/well. For high density cultures for Western blot, neurons were plated at 300k/well on poly-L-Ornithine and laminin coated 6-well plates. Neurons were infected with lentiviral particles on day in vitro (DIV) 3 or 6 and used for experiments on DIV 13-16.

### Plasmids

Plasmids for Cre (pFSW-nclcre) and dCre (pFSW-nclDeltacre) were obtained from Pascal Kaeser ^59, 114^ and possess an additional nuclear localization sequence of nucleoplasmin ^115^ in the N-terminus of EGFP to ensure optimal nuclear localization. Rescue constructs for full-length (FL) tomosyn-1 and all truncated and mutated versions of tomosyn were generated based on mouse tomosyn-1m (NP_001074813.2). All constructs contained a mScarlet-t2A sequence at N-terminal side for selection of infected neurons and were cloned into a synapsin promoter-driven construct, sequence verified and subcloned into pLenti vectors. Viral particles were produced using HEK cells ^116^.

For proteins used in single molecule study, the amino acid sequences corresponding to WT SNARE and Munc18-1 were described elsewhere in detail ^19^. The genes containing the cytoplasmic domains of rat syntaxin-1, syb2/VAMP2, mouse tomosyn-1 R-SNARE and rat Munc18-1 were cloned into the pET-SUMO vector encoding 6xHis-tag followed by a SUMO tag at the N termini. The full-length cysteine-free SNAP-25 was cloned into pET-15b vector encoding 6xHis-tag at the N terminus.

Please see the key resources table for a list of constructs used in this study.

### Protein expression, purification and SNARE preparation

All SNARE proteins and Munc18-1 were expressed in BL21 E. coli cells at 37 °C for 3 h with 1 mM isopropyl β-D-1-thiogalactopyranoside (IPTG). Proteins then were purified using Ni-NTA-agarose beads and eluted with 300 mM imidazole and exchanged to buffer containing 25 mM HEPES (pH 7.4), 140 mM KCl, and 2 mM tris(2-carboxyethyl) phosphine (TCEP). Syntaxin-1 was then biotinylated at its C-terminal Avi-tag with the biotin ligase BirA. Ternary SNARE complexes were prepared and cross-linked with DNA handles as was previously described ^19^. Briefly, syntaxin-1, SNAP-25, and syb2/VAMP2 or tomosyn were mixed at a molar ratio of 0.8:1:1.2, incubated at 4 °C, and purified using the 6xHis-tag on SNAP-25 and Ni-NTA-agarose. The eluted SNARE complexes were cross-linked with DTDP (2,2’-dithiodipyridine disulfide) treated DNA handles with a molar ratio of 50:1 in 100 mM phosphate buffer, 500 mM NaCl, pH 8.5.

### Dual-trap optical tweezers

The optical tweezers were home-built as described elsewhere. Briefly, a 1064 nm laser beam is expanded, collimated, and split into two orthogonally polarized beams, one of which is reflected by a mirror attached to a nano-positioning stage (Mad-city Labs, WI). The two beams are then combined, expanded again, and are then focused by a water-immersed 60x objective with a 1.2 numerical aperture (Olympus, PA) to form two optical traps in the sample plane in the central channel of a home-built microfluidic flow chamber. One of the two traps is stationary; the other trap can be moved using the nano-positioning stage. The outgoing laser beams are collimated by a second water-immersed objective, split again by polarization, and projected onto two position-sensitive detectors (Pacific Silicone Sensor, CA). Displacements of the trapped beads are detected by back-focal plane interferometry. Optical tweezers are remotely operated through a computer interface written in LabVIEW (National Instruments, TX). The force constants of two optical traps are determined by the Brownian motion of the trapped beads before each experiment.

### Single-molecule experiments and data analysis

An aliquot of the crosslinked protein-DNA sample was incubated with anti-digoxigenin coated polystyrene beads 2.17 µm in diameter (Spherotech, IL), diluted in phosphate-buffered saline (PBS), and injected into the top channel of a microfluidic chamber. Streptavidin-coated polystyrene beads of 1.76 µm were injected into the bottom channel. Both top and bottom channels were connected to a central channel by capillary tubes, where both kinds of beads were trapped. A single SNARE complex was tethered between two beads by bringing them close. Data were recorded at 20 kHz, mean-filtered to 10 kHz. The single-molecule experiment was conducted in PBS at 23 (±1) °C. An oxygen scavenging system was added to prevent potential protein photo-damage by optical traps. The single protein-DNA tether was pulled or relaxed by increasing or decreasing trap separation at a speed of 10 nm/s.

The methods of data analysis were described in detail elsewhere ^81, 82, 117^. Briefly, by analyzing the extension trajectories using a two-or three-state hidden-Markov modeling (HMM), the probability, extension, force, lifetime, and transition rates for each state were obtained. To relate the experimental measurements to the conformations and energy (or the energy landscape) of different SNARE states at zero force, we constructed structural models for these states based on crystal structures of the SNARE four-helix bundle and the template complex. These states were characterized by the contour lengths of the unfolded polypeptides and free energy, which were chosen as fitting parameters. The extension and energy of the whole tethered dumbbell, including the DNA handle, were calculated using the Marko-Siggia formula. Then, we computed the probability of each state based on the Boltzmann distribution and transition rates based on the Kramers’ equation. Finally, we fit the calculated state extensions, forces, probabilities, and transition rates to the corresponding experimental measurements using the nonlinear least-squares fitting, which revealed the conformations and energies of different SNARE folding states as best-fit parameters.

### Western blot

High density neuronal cultures were lysed on DIV14 in Laemmli sample buffer. Lysates were heated at 95℃ for 5 min and loaded on sodium dodecyl sulfate (SDS) polyacrylamide gels. Proteins were resolved at 100 V and transferred to nitrocellulose membranes using wet tank transfer system (Bio-Rad). Successful transfer was validated by Ponceau S staining of membranes. Membranes were blocked for 30 min in 5 % milk powder dissolved in Tris-buffered saline containing 0.1% Tween-20 detergent (TBST, pH 7.4). Thereafter, membranes were incubated overnight at 4℃ with primary antibodies dissolved in blocking buffer. The following primary antibodies were used: polyclonal rabbit anti-tomosyn (1:1000, Synaptic Systems 183103), monoclonal mouse anti-syntaxin (1:2000, Sigma S0664), monoclonal mouse anti-SNAP-25 (1:1000, Covance SMI-81R), monoclonal mouse anti-VAMP2 (1:1000, Synaptic Systems 104211), monoclonal mouse anti-actin (1:2000, Chemicon MAB1501). After the incubation with primary antibody solutions, membranes were washed three times in TBST and incubated for 1 h at room temperature with horseradish peroxidase coupled secondary antibodies (Agilent Dako) diluted at 1:10000 in blocking buffer. Membranes were rinsed three times in TBST and developed on the Odissey Fc imaging system (LI-COR Bioscience) using SuperSignal West Femto Maximum Sensitivity Substrate (Thermo Scientific). Chemiluminescent signals were analyzed using Image Studio Lite Software.

### Immunocytochemistry

Neurons were fixed with 3.7% formaldehyde (Electron Microscopy Sciences) on DIV 15. After 20 min, cells were washed with home-made phosphate-buffered saline (PBS) and stored or directly permeated using 0.5% Triton X-100 for 5 minutes. Blocking solution (BS) contained 0.2% normal goat serum and 0.1 % Triton X-100. BS was applied for 30 min followed by 2 h primary antibody incubation at room temperature (RT). Antibodies were diluted in BS. Cells were stained for MAP2 as a dendrite marker and synaptophysin-1 as a synapse marker. The following antibodies were used: polyclonal chicken anti-MAP2 (1:500, Abcam), polyclonal guinea pig anti-synaptophysin-1 (1:500, Synaptic Systems 101004), polyclonal rabbit anti-tomosyn (1:500, Synaptic Systems 183103). After washing 3 times with PBS, cells were incubated with secondary antibodies diluted 1:1000 in BS for 1 h. The following secondary antibodies were used: goat anti-chicken Alexa546, goat anti-guinea pig Alexa488, goat anti-rabbit Alexa647 (Molecular Probes). After another 3 washes in PBS, coverslips were mounted on microscope slides using DABCO-Mowiol (Invitrogen). Cells were imaged on a confocal laser-scanning microscope (Nikon Eclipse Ti A1) using a 40x oil immersion objective (NA 1.3). Only EGFP-positive autapses were imaged. Neuronal morphology, synapse numbers and the average tomosyn intensity per neuron were analysed using the semi-automated platform SynD ^65^ (www.johanneshjorth.se/SynD).

### Electrophysiology

Autaptic neurons were subjected to whole-cell voltage-clamp recordings (Vm = −70 mV) on DIV 13-16. Experiments were performed at room temperature with borosilicate glass pipettes (Science products GmbH, 2.5-4.5 Mohm) filled with (in mM): 136 KCl, 1 HEPES, 0.6 MgCl2*6H2O, 4 ATP-Mg, 0.3 GTP-Na, 12 phosphocreatine dipotassiumsalt and 50 U/ml phosphocreatine kinase (pH = 7.3, ∼300 mOsmol). External solution (aCSF) contained the following (in mM): 10 HEPES, 10 Glucose, 140 NaCl, 2.4 KCl, 4 MgCl2 and 2 CaCl2 (pH = 7.30, ∼300 mOsmol). Solutions were made from stock with HEPES and Glucose added fresh, solutions were filter-sterilized and stored at 4°C until use. Patch-clamp recordings were performed with a MultiClamp 700B amplifier and Digidata 1550B or an Axopatch 200B amplifier and Digidata 1440A, controlled by Clampex 10.6 software (Molecular Devices). For gap-free recordings of spontaneous miniature EPSCs, the sampling rate was set to 20 kHz and low-pass Bessel filter was set to 5-6 kHz. For episodic stimulations, the sampling rate was set to 10 kHz and low-pass Bessel filter to 2 kHz. Resistance was compensated by 70-80% (bandwidth 7.52 Hz). Feedback resistor 500 Mohm was adjusted only if EPSCs were larger than 20 nA. Action potentials were elicited by a 1 ms depolarization to 30 mV. Recordings were excluded if series resistance was higher than 15 Mohm or leak current was larger than 300 pA. GABAergic recordings were identified based on their postsynaptic decay kinetics and excluded. Evoked responses smaller than 500 pA were only included for EPSC size, not plasticity measurements. Offline analysis was performed with MATLAB R2018b and R2019a (Mathworks) using custom-written software routines (github.com/vhuson/viewEPSC). EPSC kinetics were calculated in Clampfit 10.6.

### Hypertonic sucrose application

250 mM and 500 mM sucrose (Sigma-Aldrich) solutions were freshly made from aCSF at the start of each experimental week. Sucrose was applied via gravity infusion through a custom-made barrel system controlled by a perfusion Fast-Step delivery system (SF-77B, Warner instrument corporation). Speed of flow was controlled with an Exadrop precision flow rate regulator (B Braun). Multiple sucrose applications were performed in the following order: 250 mM, 3 minutes recovery time, 500 mM, 30 seconds recovery time, 500 mM. Each application of sucrose lasted 7 seconds. In between sucrose applications, cells were constantly perfused with aCSF. In manual analysis, the charge released within the first 4 seconds of sucrose application was calculated. Alternatively, the sucrose traces were fitted with a minimal vesicle state model as published previously ^72, 73^ using MATLAB code kindly shared by Vincent Huson. This method yields a more accurate measure of the readily-releasable pool (RRP) compared to other methods by accounting for ongoing priming. Additional measures obtained from this fitting are priming and de-priming rates, the fusion rate (k2) and the energy barrier for fusion (ΔEa). The release rate and energy barrier at 0 mM sucrose were obtained by analyzing the mEPSCs within the first 6 seconds before the first application of sucrose, recorded in the same traces. The frequency of miniature EPSCs (mEPSCs) within this time was multiplied by the average charge of one mEPSC; this value was then divided by the RRP measure to calculate k2 at 0 mM sucrose. An estimation of the energy barrier results from computing the natural logarithm of k2 (see ^72^ for method details). The average estimation from WT cells was used to calculate the change in the energy barrier evoked by applications of sucrose. The time of sucrose onset was analyzed using Clampfit 10.6 (Molecular Devices) by documenting the position of a cursor manually placed at the start of a sucrose response.

### Electron Microscopy

Dissociated hippocampal mouse neurons (5 k/well) were plated on pre-grown cultures of rat glia on sapphire disks (Wohlwend GmbH) to form micro-networks of 2–10 neurons per sapphire disk. Prior to culturing, sapphire disks were etched for 30 minutes in 60% sulfuric acid, washed, incubated in 3M KOH overnight, washed and dried before carbon coating and subsequent baking at 180 C for 2 hours. Sapphire discs were coated by a mixture of 0.1 mg/ml poly-d-lysine (Sigma), 0.7 mg/ml rat tail collagen (BD Biosciences), and 10 mM acetic acid (Sigma) and placed in an agarose-coated 12-well plate to form glia monolayer islands selectively on sapphire disks. The sapphire disks were cryofixed in an EM-PACT2 (Leica Microsystems) high-pressure freezer in 5% trehalose/10% BSA in 0.05M phosphate buffer (pH 7.4, 320 mOsm) cryoprotectant. Frozen samples were postfixed in 1% OsO4/ 5% saturated K4Fe(CN)6 in H2O in acetone at −90°C for 74 hours and brought to 0°C at 5°/h. After several washes with ice-cold acetone, the sapphire disks were washed with propylene oxide and infiltrated by an increasing Epon concentration series. The samples were embedded in fresh Epon overnight and left to polymerize at 65°C for 48 h. Sapphire disks were separated from the Epon by dipping the samples in boiling water and liquid nitrogen and regions with micro-networks were selected by light microscopy. These regions were cut out and mounted on pre-polymerized Epon blocks for ultrathin sectioning. Ultrathin sections (80 nm) were cut parallel to the sapphire disk, collected on single-slot, formvar-coated copper grids, and stained in uranyl acetate and lead citrate. Hippocampal synapses were randomly selected at low magnification using an electron microscope (JEOL1010) while being blinded for the experimental conditions. For each condition, the number of docked SVs, total SV number, and active zone length were measured on digital images taken at 80,000-fold magnification using custom-written semiautomatic image analysis software running in Matlab (Mathworks). For all morphological analyses the following requirements were set: clearly recognizable synapses with intact synaptic plasma membranes with a recognizable pre-and postsynaptic area and defined SV membranes. SVs were defined as docked if there was no distance visible between the SV membrane and the active zone membrane.

### Statistics

Datasets on single neuron parameters consist of several neuronal cultures (N = number of independent cultures), in which different coverslips from the same culture are infected with different viruses resulting in separate experimental groups (e.g. control and cDKO correspond to deltaCre-EGFP and Cre-EGFP infected neurons), from which multiple observations (n = individual neurons) are taken. To account for the nested nature of our data, we performed fixed linear regression in which culture was included as a linear predictor. Outliers, defined as datapoints more than 3 standard deviations above or below the group mean, were removed. Data were then standardized into Z-scores by grand mean centering. A fixed linear regression model was fitted to the standardized data using the lm() function in R (version 4.1.0). A one-way anova (analysis of variance) was used to assess whether including the experimental group as a second linear predictor (formula = y ∼ Group + Culture) statistically improved the fit of a model without group information (formula = y ∼ 1 + Culture). When more than two experimental groups were present, posthoc analysis was performed repeating the fixed linear regression on pair-wise subsets of the data, and p-value thresholds were Bonferroni-adjusted to account for multiple testing. Electron microscopy data on low density networks was tested by multilevel analysis using the lme() function in R (version 4.1.0) with sapphire disc as nesting level with the highest intracluster correlation of the data. A one-way anova was used to assess whether including the experimental group as a predicted variable (formula = y ∼ Group, random=∼1|Sapphire, method = “ML”) statistically improved the fit of a model without group information (formula = y ∼ 1, random=∼1|Sapphire, method = “ML”). Post hoc pairwise contrasts were extracted by estimated marginal means (emmeans() function), and p-value thresholds were Bonferroni-adjusted to account for multiple testing. Detailed information per dataset (mean, SEM, n and statistical details) is shown in supplemental item 1.

---

KEY RESOURCES TABLE

KEY RESOURCES TABLE
| REAGENT or RESOURCE | SOURCE | IDENTIFIER |
| --- | --- | --- |
| **Antibodies** |  |  |
| polyclonal chicken anti-MAP2 | Abcam | Cat#ab5392; RRID: AB_2138153 |
| polyclonal guinea pig anti-synaptophysin-1 | Synaptic Systems | Cat#101004; RRID: AB_1210382 |
| polyclonal rabbit anti-tomosyn-1 | Synaptic Systems | Cat#183103; RRID: AB_2619878 |
| monoclonal mouse anti-VAMP2 | Synaptic Systems | Cat#104211; RRID: |
| monoclonal mouse anti-SNAP25 | Covance | Cat# SMI-81R |
| monoclonal mouse anti-syntaxin | Sigma | Cat# S0664; RRID: |
| monoclonal mouse anti-actin | Chemicon | Cat# MAB1501 |
| **Experimental models: Organisms/strains** |  |  |
| Mouse: Tom1<sup>lox</sup>; Strain C57BL/6 | This paper |  |
| Mouse: Tom2<sup>lox</sup>; Strain C57BL/6 | Geerts et al.,<sup>47</sup> |  |
| Rat: Strain Wistar (CrI:WI) | Charles River | Strain code: 003 |
| **Recombinant DNA** |  |  |
| Plasmid: pFSW-nclcre | Kaeser et al.,<sup>59</sup> |  |
| Plasmid: pFSW-nclDeltacre | Kaeser et al.,<sup>59</sup> |  |
| Plasmid: pSyn(pr)mScarlet-T2A-tomosyn1 | This paper | NP_001074813.2 |
| Plasmid: pSyn(pr)mScarlet-T2A-tomosyn1(F1097A) | This paper |  |
| Plasmid: pSyn(pr)mScarlet-T2A-tomosyn1(Delta aa1105-1116) | This paper |  |
| Plasmid: pSyn(pr)mScarlet-T2A-tomosyn1(aa1-1047) | This paper |  |
| Plasmid: pSyn(pr)mScarlet-T2A-tomosyn1(aa1-994) | This paper |  |
| Plasmid: pSyn(pr)mScarlet-T2A-tomosyn1(aa1-1047)-VAMP2(aa28-96) | This paper |  |
| Plasmid: pSyn(pr)mScarlet-T2A-tomosyn1(aa1-1047)-VAMP2(aa28-84)-tomosyn1(aa1105-1116) | This paper |  |
| Plasmid: pSyn(pr)mScarlet-T2A-tomosyn1(aa1-1104)-VAMP2(aa85-96) | This paper |  |
| pET-SUMO-syntaxin(aa1-265) | Jiao et al.,<sup>19</sup> |  |
| pET-SUMO-VAMP2(aa1-96) | Jiao et al.,<sup>19</sup> |  |
| pET-SUMO-tomosyn(aa1045-1116) | This paper | NP_001074813.2 |
| pET-15b-SNAP-25(aa1-206) | Jiao et al.,<sup>19</sup> |  |
| pET-SUMO-Munc18-1(aa1-594) | Jiao et al.,<sup>19</sup> |  |
| **Software and Algorithms** |  |  |
| Matlab R2019a | Mathworks |  |
| R version 4.1.0 | R Core Team, 2021 |  |
| Clampex 10.6 | Molecular Devices |  |
| SynD | Schmitz et al., 2011 |  |
| Image Studio Lite Software | LI-COR Biosciences |  |

