## Supplemental figures for "Tomosyns attenuate SNARE assembly and synaptic depression by binding to VAMP2-containing template complexes"

**Figure S1**

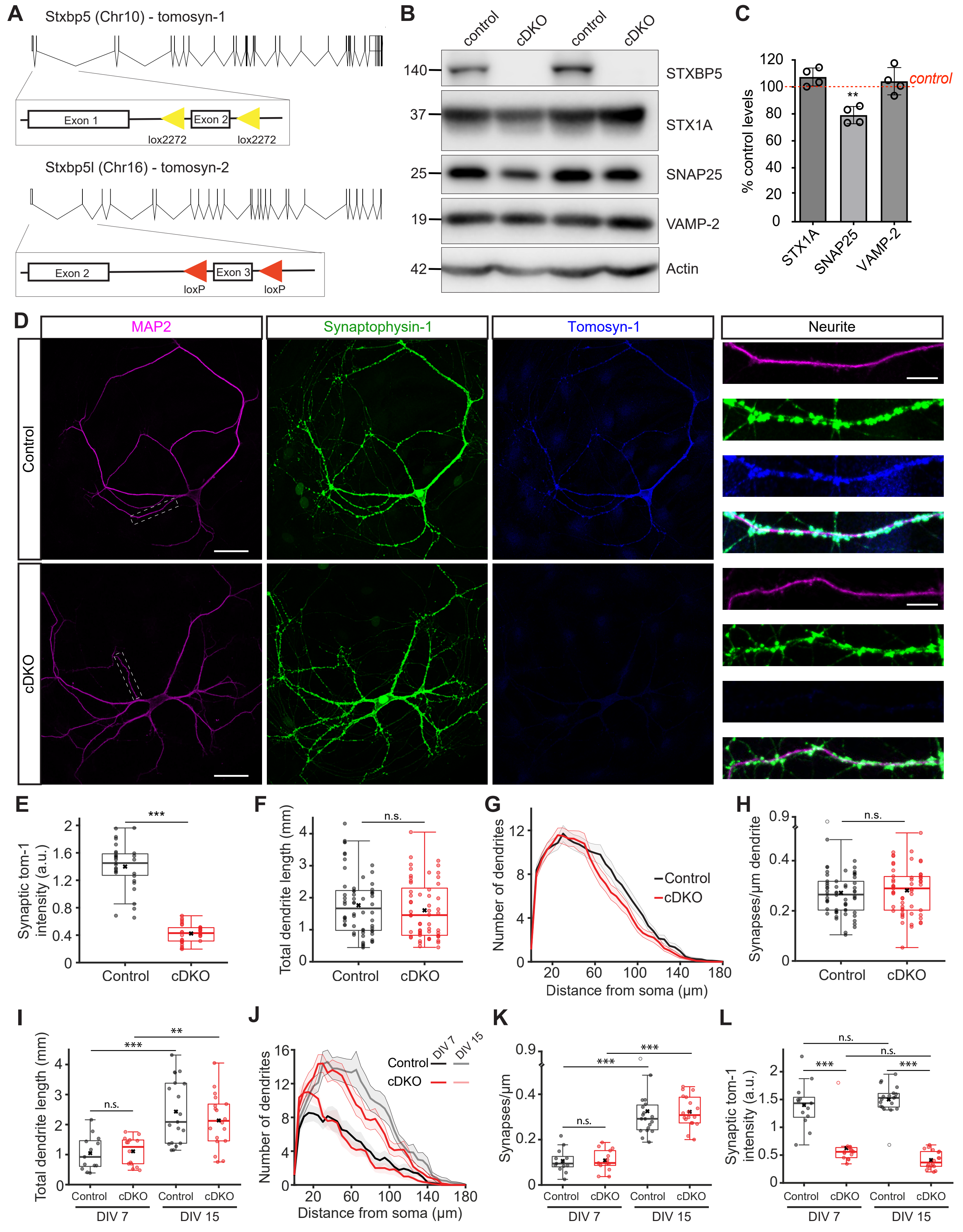

A
Stxbp5 (Chr10) - tomosyn-1
Exon 1
Exon 2
lox2272
lox2272
Stxbp5l (Chr16) - tomosyn-2
Exon 2
Exon 3
loxP
loxP
B
control
cDKO
control
cDKO
140
37
25
19
42
STXBP5
STX1A
SNAP25
VAMP-2
Actin
C
% control levels
control
STX1A
SNAP25
VAMP-2
D
MAP2
Synaptophysin-1
Tomosyn-1
Neurite
Control
cDKO
E
Synaptic tom-1 intensity (a.u.)
Control
cDKO
F
Total dendrite length (mm)
Control
cDKO
G
Number of dendrites
Control
cDKO
Distance from soma (µm)
H
Synapses/µm dendrite
Control
cDKO
I
Total dendrite length (mm)
Control
cDKO
DIV 7
DIV 15
J
Number of dendrites
Control
cDKO
DIV 7
DIV 15
Distance from soma (µm)
K
Synapses/µm
Control
cDKO
DIV 7
DIV 15
L
Synaptic tom-1 intensity (a.u.)
Control
cDKO
DIV 7
DIV 15

**Figure S2**

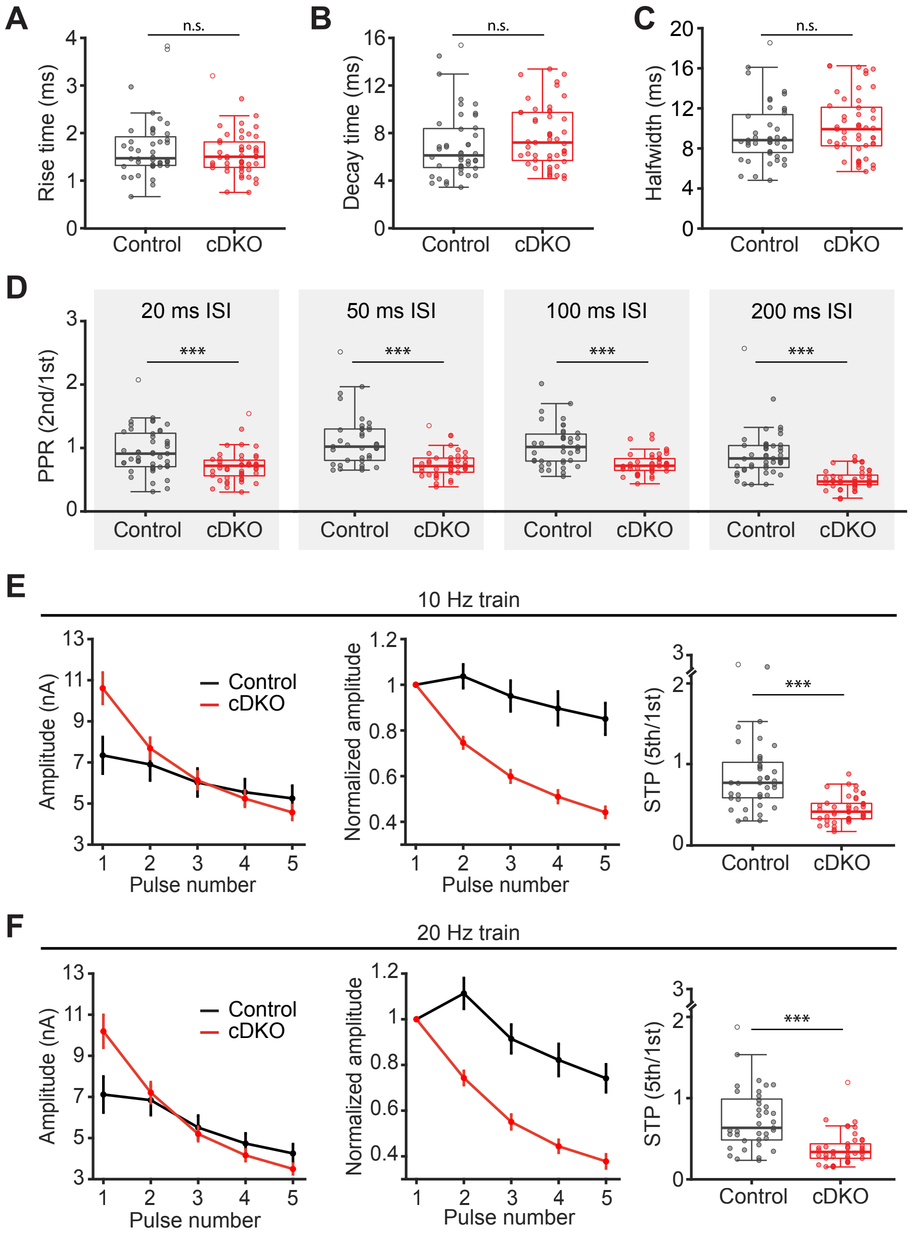
A
B
C
n.s.
Rise time (ms)
Decay time (ms)
Halfwidth (ms)
Control
cDKO
D
20 ms ISI
50 ms ISI
100 ms ISI
200 ms ISI
***
PPR (2nd/1st)
E
10 Hz train
Amplitude (nA)
Normalized amplitude
STP (5th/1st)
Pulse number
F
20 Hz train

Figure S3

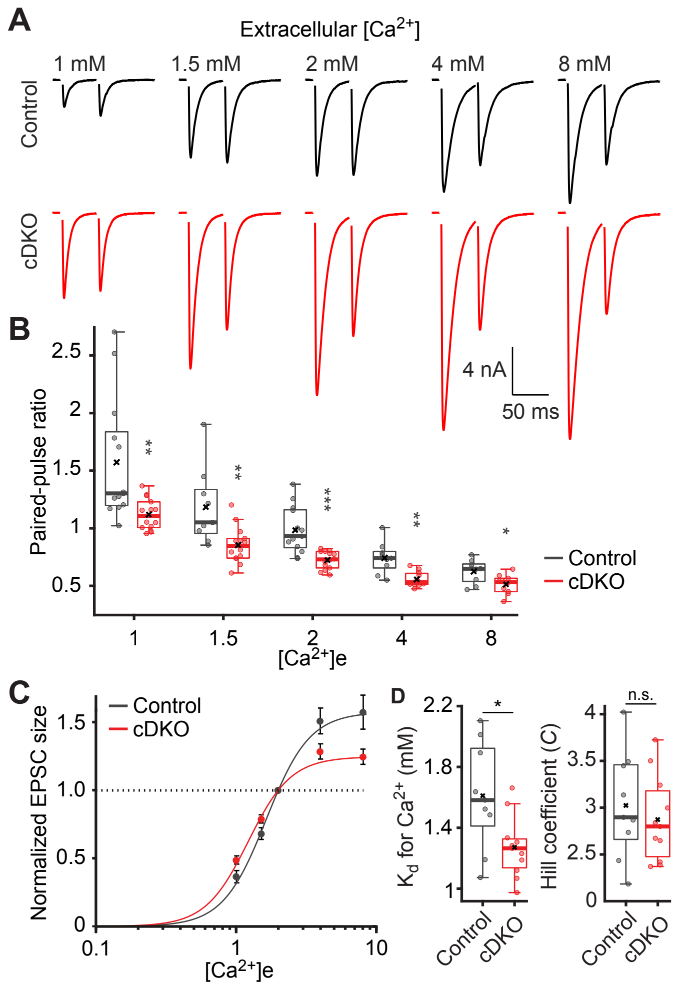
A
Extracellular [Ca2+]
1 mM
1.5 mM
2 mM
4 mM
8 mM
Control
cDKO
4 nA
50 ms
B
Paired-pulse ratio
2.5
2
1.5
1
0.5
**
**
***
**
*
Control
cDKO
1
1.5
2
4
8
[Ca2+]e
C
Normalized EPSC size
1.5
1.0
0.5
0
0.1
1
10
[Ca2+]e
Control
cDKO
D
Kd for Ca2+ (mM)
2.2
1.8
1.4
1
*
Control
cDKO
Hill coefficient (C)
4
3.5
3
2.5
0
n.s.
Control
cDKO

**Figure S4**

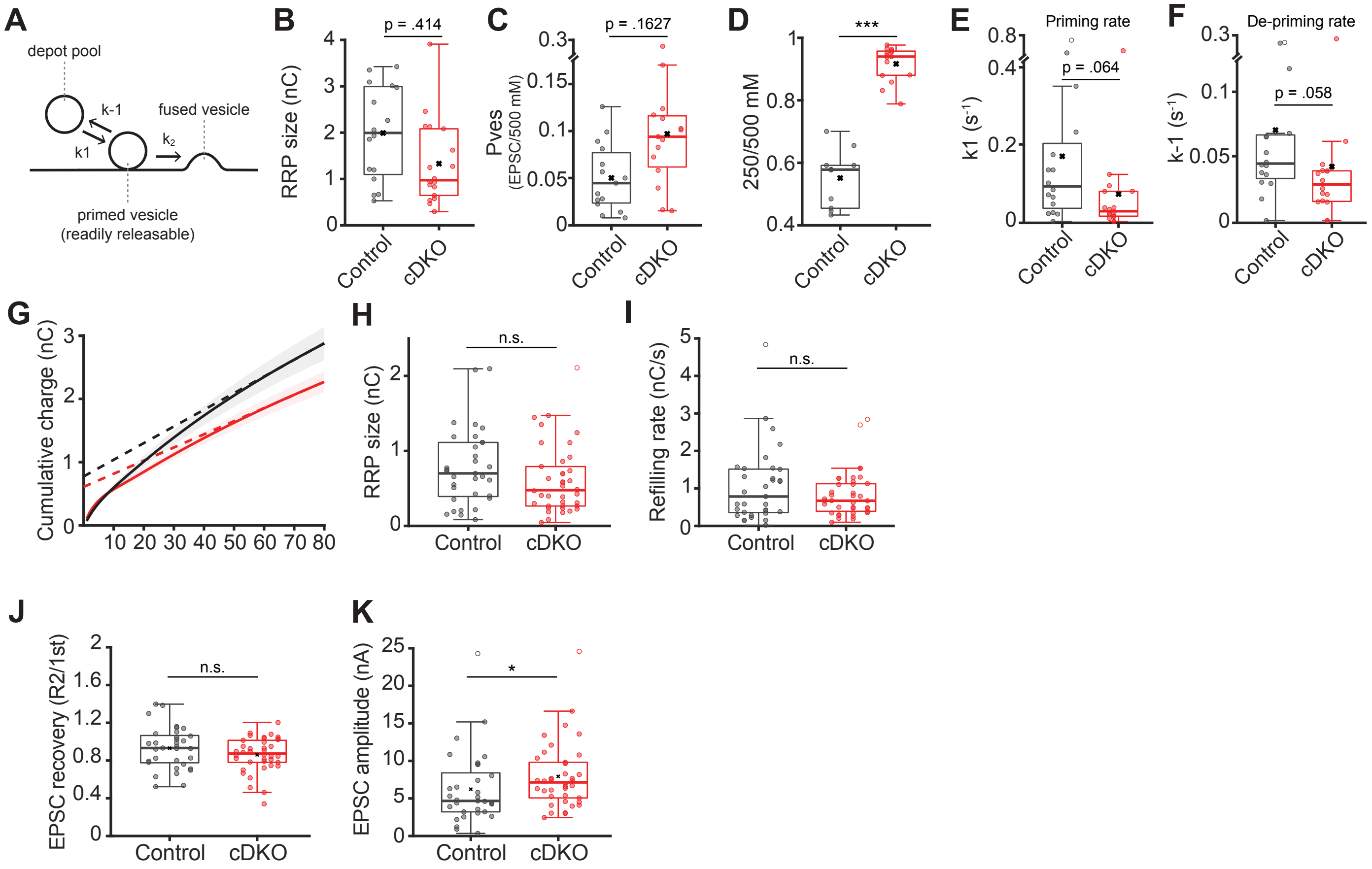
A
depot pool
k-1
k1
fused vesicle
k2
primed vesicle
(readily releasable)
B
p = .414
RRP size (nC)
Control
cDKO
C
p = .1627
Pves
(EPSC/500 mM)
Control
cDKO
D
***
250/500 mM
Control
cDKO
E
Priming rate
p = .064
k1 (s-1)
Control
cDKO
F
De-priming rate
p = .058
k-1 (s-1)
Control
cDKO
G
Cumulative charge (nC)
H
n.s.
RRP size (nC)
Control
cDKO
I
n.s.
Refilling rate (nC/s)
Control
cDKO
J
n.s.
EPSC recovery (R2/1st)
Control
cDKO
K
*
EPSC amplitude (nA)
Control
cDKO

**Figure S5**

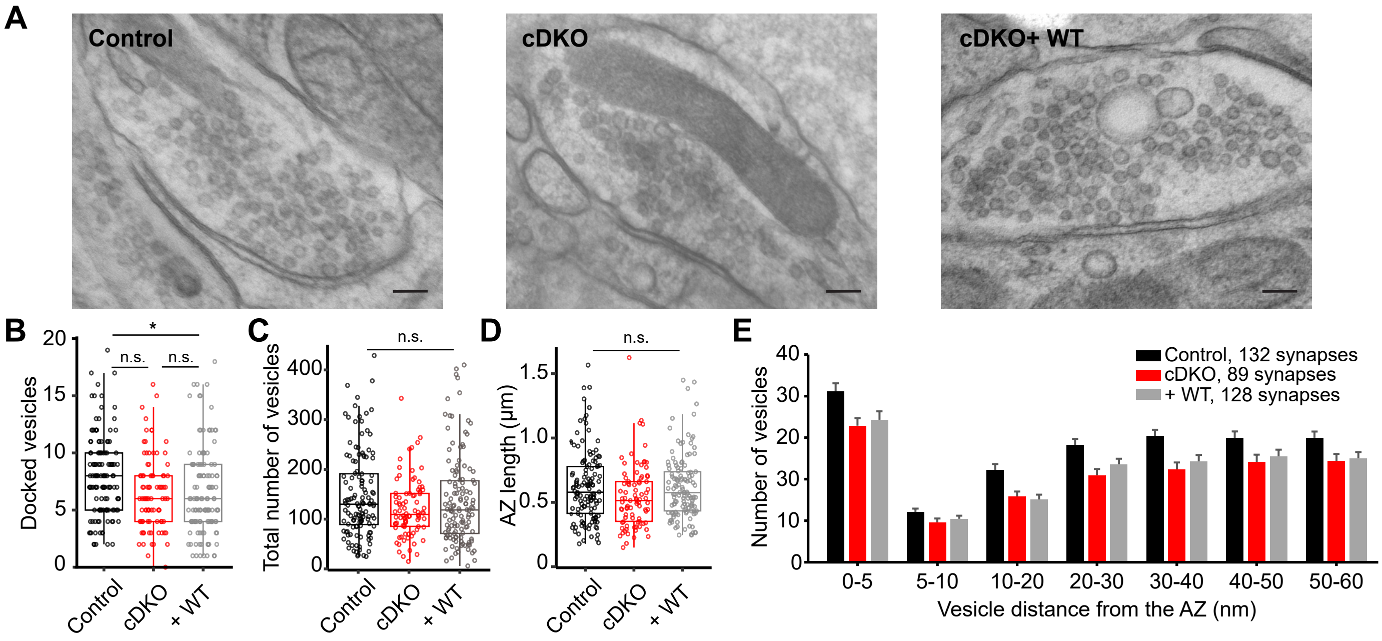
A
Control
cDKO
cDKO+ WT
B
Docked vesicles
*
n.s.
n.s.
Control
cDKO
+ WT
C
Total number of vesicles
n.s.
D
AZ length (µm)
n.s.
E
Number of vesicles
Control, 132 synapses
cDKO, 89 synapses
+ WT, 128 synapses
0-5
5-10
10-20
20-30
30-40
40-50
50-60
Vesicle distance from the AZ (nm)

**Figure S6**

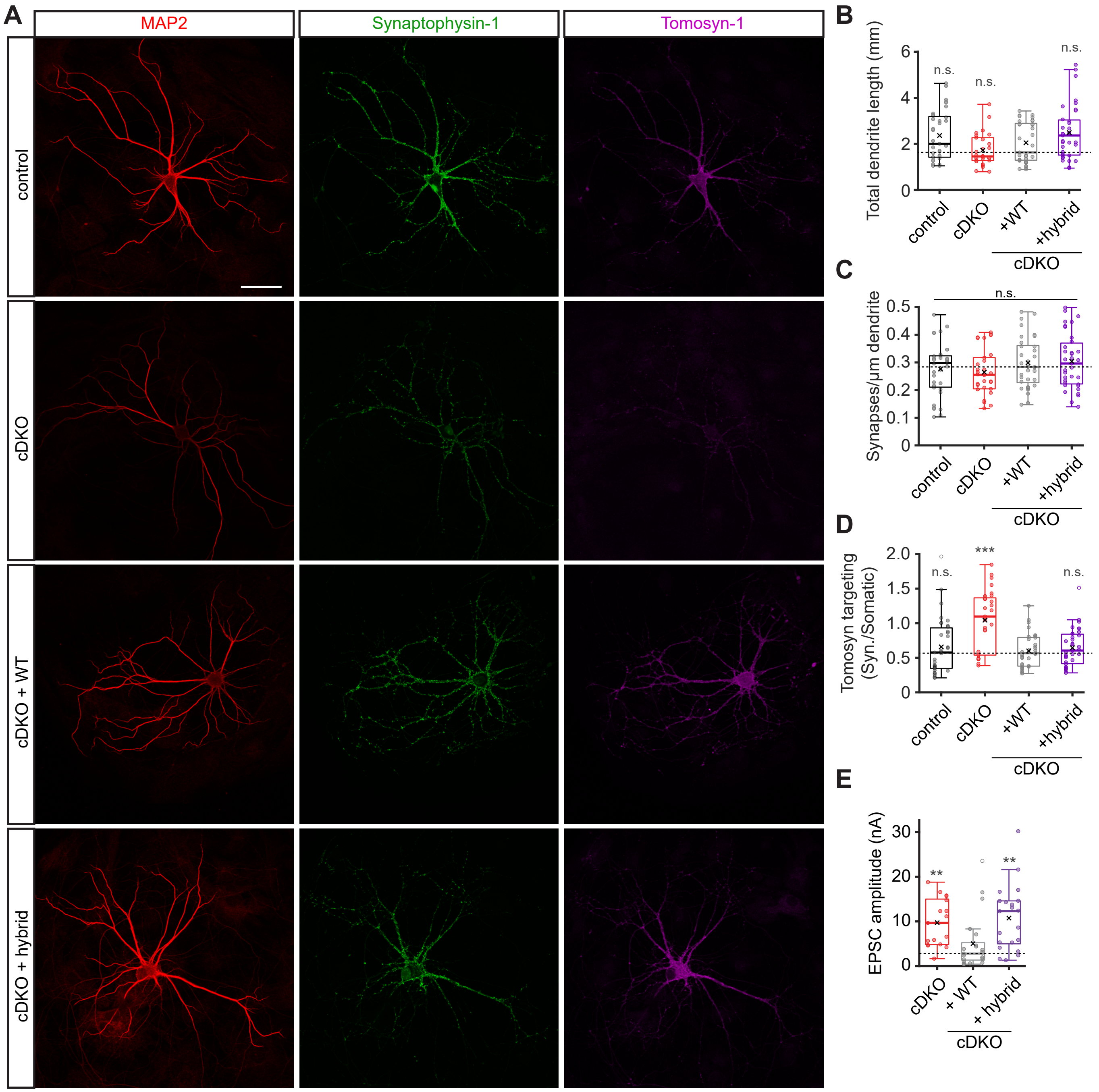
A
MAP2
Synaptophysin-1
Tomosyn-1
control
cDKO
cDKO + WT
cDKO + hybrid
B
Total dendrite length (mm)
n.s.
n.s.
n.s.
control
cDKO
+WT
+hybrid
cDKO
C
Synapses/µm dendrite
n.s.
control
cDKO
+WT
+hybrid
cDKO
D
Tomosyn targeting (Syn./Somatic)
n.s.
***
n.s.
control
cDKO
+WT
+hybrid
cDKO
E
EPSC amplitude (nA)
**
**
cDKO
+ WT
+ hybrid
cDKO

**Figure S7**

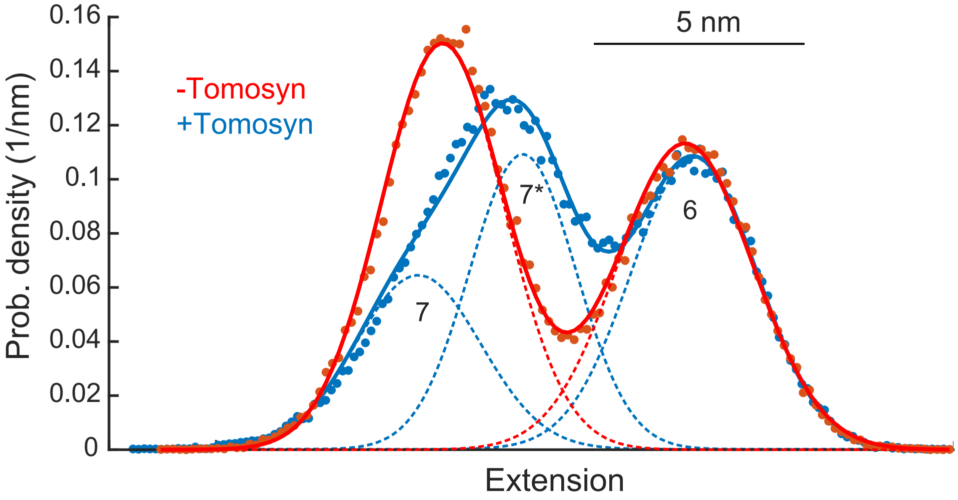

0.16
0.14
0.12
0.1
0.08
0.06
0.04
0.02
0
Prob. density (1/nm)
Extension
5 nm
-Tomosyn
+Tomosyn
7
7*
6

**Figure S8**

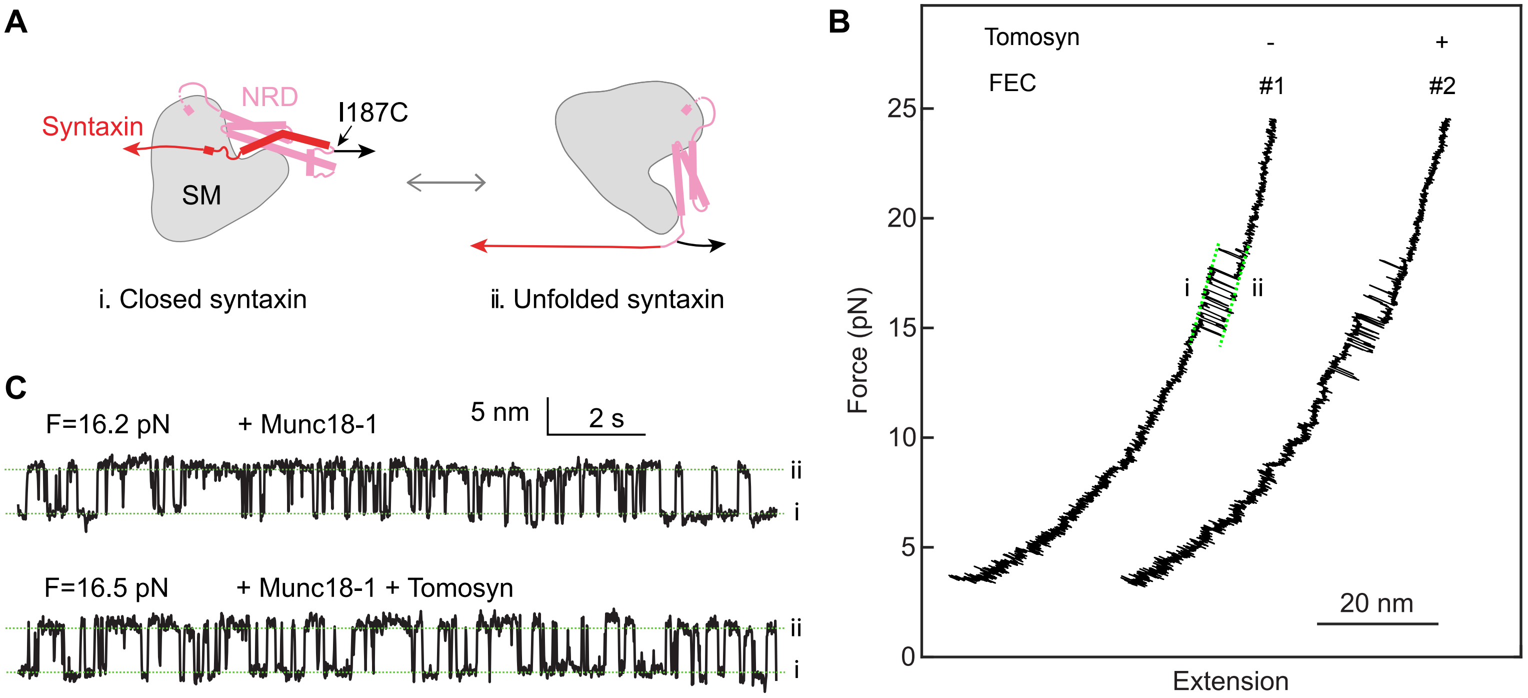
A
Syntaxin
NRD
I187C
SM
i. Closed syntaxin
ii. Unfolded syntaxin
B
Tomosyn
-
+
FEC
#1
#2
Force (pN)
25
20
15
10
5
0
i
ii
20 nm
Extension
C
F=16.2 pN
+ Munc18-1
5 nm
2 s
ii
i
F=16.5 pN
+ Munc18-1 + Tomosyn
ii
i

**Figure S9**

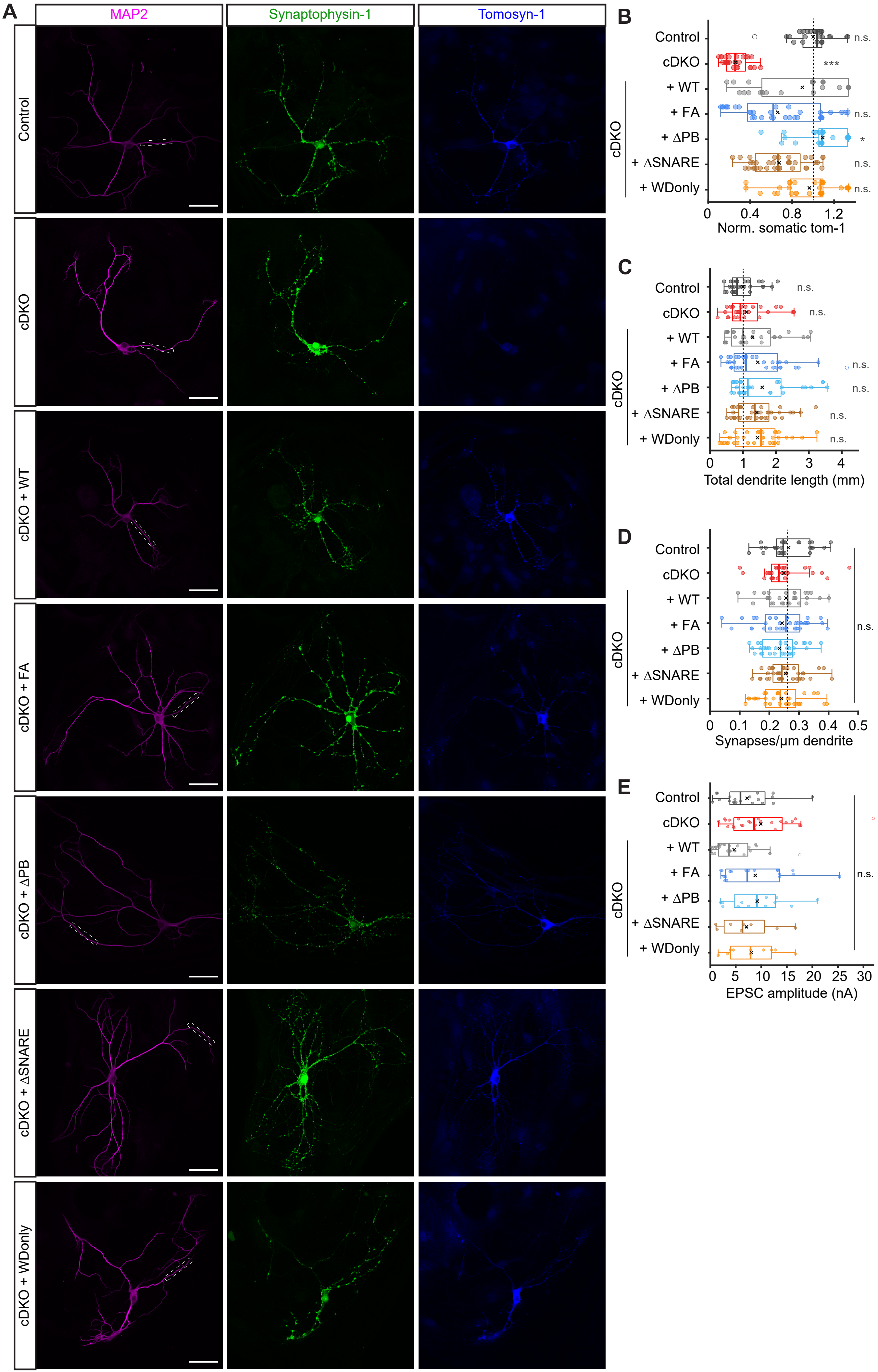

A
MAP2
Synaptophysin-1
Tomosyn-1
Control
cDKO
cDKO + WT
cDKO + FA
cDKO + ΔPB
cDKO + ΔSNARE
cDKO + WDonly
B
Control
cDKO
+ WT
+ FA
+ ΔPB
+ ΔSNARE
+ WDonly
cDKO
n.s.
***
n.s.
*
n.s.
n.s.
0
0.4
0.8
1.2
Norm. somatic tom-1
C
Control
cDKO
+ WT
+ FA
+ ΔPB
+ ΔSNARE
+ WDonly
cDKO
n.s.
n.s.
n.s.
n.s.
n.s.
n.s.
0
1
2
3
4
Total dendrite length (mm)
D
Control
cDKO
+ WT
+ FA
+ ΔPB
+ ΔSNARE
+ WDonly
cDKO
n.s.
0
0.1
0.2
0.3
0.4
0.5
Synapses/µm dendrite
E
Control
cDKO
+ WT
+ FA
+ ΔPB
+ ΔSNARE
+ WDonly
cDKO
n.s.
0
5
10
15
20
25
30
EPSC amplitude (nA)

**Figure S10**

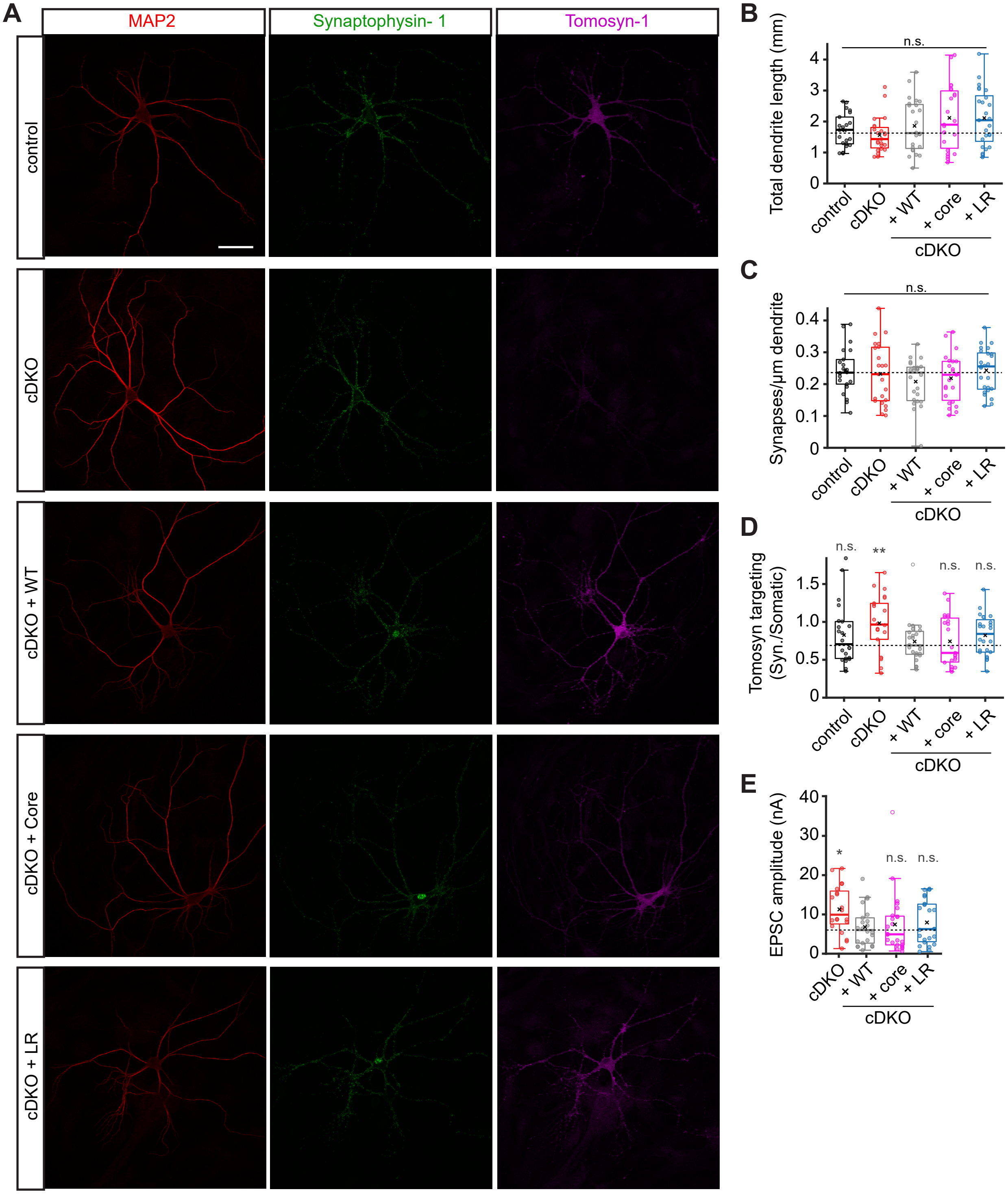

A
MAP2
Synaptophysin- 1
Tomosyn-1
control
cDKO
cDKO + WT
cDKO + Core
cDKO + LR
B
Total dendrite length (mm)
n.s.
0
1
2
3
4
control
cDKO
+ WT
+ core
+ LR
cDKO
C
Synapses/μm dendrite
n.s.
0
0.1
0.2
0.3
0.4
control
cDKO
+ WT
+ core
+ LR
cDKO
D
Tomosyn targeting (Syn./Somatic)
n.s.
**
n.s.
n.s.
0
0.5
1.0
1.5
control
cDKO
+ WT
+ core
+ LR
cDKO
E
EPSC amplitude (nA)
*
n.s.
n.s.
0
10
20
30
40
cDKO
+ WT
+ core
+ LR
cDKO

### Supplemental figure legends

**Figure S1: A novel conditional KO mouse for both tomosyn paralogs.**
(**A**) Structures of the tomosyn-1 and tomosyn-2 genes. Zoom-ins to early exons show floxed regions around exon 2 of tomosyn-1 and exon 3 of tomosyn-2.
(**B**) Example Western blot showing loss of tomosyn-1 (STXBP5) and levels of synaptic SNAREs in cDKO high-density neuronal cultures. Numbers on the left indicate approximate molecular weight of the detected bands in kDa.
(**C**) Quantification of synaptic SNAREs expression from Western blots exemplified in **B**. cDKO levels were normalized to control levels in the corresponding culture. Four independent cultures were analyzed. Data are shown as mean ± SD and were analyzed using one sample t-test. **p<.01.
(**D**) Example images of autaptic hippocampal neurons immunostained for MAP2 as a dendrite marker, synaptophysin-1 as a synapse marker, and tomosyn-1 (scale bar = 50 µm). Dashed boxes in MAP2 images correspond to dendrite zoom-ins on the right. Dendrite zoom-ins show synaptic localization of tomosyn-1 and loss of expression in cDKO neurons (scale bar = 10 µm).
(**E-H**) Quantifications from images shown in **D.** (**E**) Loss of tomosyn expression in synapses was confirmed by measuring the intensity of the tomosyn-1 signal within synaptophysin-1 puncta in cDKO (n = 30/2) and control (n = 31/2) neurons. ***p<.001 (**F**) Total dendritic length was determined by tracing of the MAP2 signal. Control n = 62/4, cDKO n = 57/4. (**G**) Sholl plot showing similar dendritic complexity of both genotypes. (**H**) Synapse density was determined by the number of synaptophysin-1-positive puncta within one µm dendrite. Control n = 62/4, cDKO n = 57/4.
(**I-K**) Morphological analysis from immunostainings at day in vitro (DIV) 7 and 15 shows similar morphology of both groups throughout development. (**I**) Total dendritic length derived from MAP2 tracing. **p<.0033, ***p<.00033 (**J**) Sholl plot of dendrites. (**K**) Synapse density was determined from synaptophysin-1-positive puncta within one µm dendrite. ***p<.00033
(**l**) Signal intensity of tomosyn-1 in synaptophysin-positive puncta confirms loss of tomosyn at both DIV7 and DIV15. DIV7: control n = 13/1, cDKO n = 14/1; DIV15: control n = 19/1, cDKO n = 18/1. ***p<.00033
In **G** and **J**, data are presented as mean ± SEM. P-values were determined by ANOVA and Bonferroni-corrected if applicable. Abbreviations: n.s. (not significant); DIV (day in vitro).

**Figure S2: Normal EPSC kinetics and more quantifications of release probability and short-term plasticity.**
(**A-C**) Analysis of the kinetics of single EPSCs. Control n = 44/6, cDKO n = 49/6. (**A**) 20 – 80% rise time. (**B**) 100 – 50% decay time. (**C**) EPSC width at half maximum.
(**D**) Paired-pulse ratios at different inter-pulse intervals. Same data as in Figure 1 G. Control n = 37 – 47/6, cDKO n = 42 – 47/6.
(**E**) Same analysis as in figure 2 H-K but for a 10 Hz train of 5 pulses. The rundown of the absolute (left) and normalized (center) amplitude was plotted and the ratio of the fifth over the first amplitude was calculated (right) to quantify short-term plasticity. Control n = 39/6, cDKO n =44/6.
(**F**) Same as in **E** but for a 20 Hz train. Control n = 39/6, cDKO n = 42/6.
In left and center plots in **E** and **F**, data are presented as mean ± SEM. ***p<.001 as determined by ANOVA. Abbreviations: n.s. (not significant).

**Figure S3: Tomosyns increase the calcium affinity of the release machinery.**

(**A-B**) Autaptic hippocampal neurons were stimulated with paired pulses of 50 ms intervals at different extracellular calcium concentrations. Control n = 9 – 13/4, cDKO n = 10 – 14/4. (**A**) Example traces. (**B**) The paired-pulse ratio (PPR) was calculated by dividing the amplitude of the second pulse by the amplitude of the first pulse.
(**C**) Concentration-response curves of the normalized EPSC amplitudes from paired pulses shown in **A-B**. Amplitudes were normalized to responses in standard 2 mM $[Ca^{2+}]_e$.
(**D**) A Hill function was fitted to the data shown in **C** from which the dissociation constant (Kd) for calcium (left) and the Hill coefficient (right) were calculated. Control n = 9/4, cDKO n = 11/4.
In **C**, data are presented as mean ± SEM. *p<.05, **p<.01, ***p<.001 as determined by ANOVA. Abbreviations: n.s. (not significant).

**Figure S4: Fitting of an energy barrier model yields similar results as manual analysis of sucrose traces.**
(**A**) Illustration of the minimal vesicle state model. Vesicles first start out in a depot pool of vesicles from which they transition to a primed state at a priming rate k1. De-priming occurs at a rate of k-1. Primed vesicles will fuse with a rate k2.
(**B**) RRP size as estimated from the fitted data. Control n = 16/6, cDKO n = 18/6.
(**C**) Pves calculated as the ratio of the charge released by a single EPSC to the fitted RRP. Control n = 15/6, cDKO n = 15/6.
(**D**) The fraction of the RRP released by the submaximal sucrose concentration as calculated from the fitted data. Control n = 10/6, cDKO n = 15/6.
(**E**) Priming and (**F**) de-priming rates during the 500 mM sucrose application, derived from the fitting. Control n = 16/6, cDKO n = 16/6.
(**G**) Neurons were stimulated with 80 action potentials at 40 Hz (see Figure 2 G). The cumulative charge released during the train was plotted and a line was drawn through the last 20 pulses to back-extrapolate to the y-intercept, which marks the estimate for the RRP (readily releasable pool) size.
(**H**) RRP size as calculated by back-extrapolation shown in **G**. Control n = 33/6, cDKO n = 38/6.
(**I**) The refilling rate of vesicles during the pulse is estimated from the slope of the back-extrapolation line shown in **G**.
(**J**) The amplitude of recovery pulse R2 (see Figure 2 G) was divided by the first amplitude of the train. (**K**) The absolute amplitude of recovery pulse R2. Control n = 31/6, cDKO n = 35/6
In **G**, data are presented as mean ± SEM (shaded area). ***p<.001 as determined by ANOVA.

**Figure S5: Loss of tomosyns does not increase vesicle docking.**
(**A-E**) High-pressure freeze electron microscopy was performed on control (n = 132 synapses), cDKO (n = 89 synapses), and cDKO neurons expressing tomosyn-1m (+ WT; n = 128 synapses). (**A**) Example images. Scale bar = 100 nm. (**B**) No significant difference between control and cDKO synapses is detected, but the number of docked vesicles is reduced in + WT synapses compared to control. (**C**) The total number of synaptic vesicles and (**D**) the length of the active zone in the cross section is normal in all groups. (**D)** The number of vesicles within distinct increments from the active zone.

Bonferroni-corrected alpha-levels: *p<.01667. Abbreviations: n.s. (not significant).

**Figure S6: The tomosyn-VAMP2 hybrid is properly targeted to synapses.**

(**A-D**) Morphological analysis of neurons infected with wild-type tomosyn or hybrid tomosyn (see Figure 3). Control n = 35/3, cDKO n = 30/3, + WT n = 33/3, + Hybrid n = 35/3. (**A**) Example images of autaptic hippocampal neurons immunostained for MAP2, synaptophysin-1, and tomosyn-1. Scale bar = 50 µm. (**B**) Total dendrite length as derived from the MAP2 mask. (**C**) Synapse density as derived from the synaptophysin-1-positive puncta within the MAP2 mask. (**D**) The ratio of the tomosyn-1 signal intensity in synapses to the signal in the soma was calculated to demonstrate synaptic targeting.
(**E**) Absolute EPSC amplitudes.
Bonferroni-corrected alpha-levels: *p<.0166, ***p<.00033; panel E: **p<0.005. Grey asterisks show comparison to + WT group. Abbreviations: n.s. (not significant).

**Figure S7. The probability density functions (PDFs) of the time-dependent extensions at constant forces revealed an intermediate state induced by the tomosyn SNARE motif (state 7*).**
The PDFs can be well fitted by a sum of two Gaussian functions in the absence of the tomosyn SNARE motif and three Gaussian functions when present. The three peaks represent the template complex (state 7), the tomosyn-bound template complex (state 7*), and the Munc18-1-bound open syntaxin (Figure 5C).

**Figure S8. The tomosyn SNARE motif does not affect the folding of Munc18-1-bound closed syntaxin-1**
(**A**) Schematic diagrams of Munc18-1-bound closed syntaxin (i) and unfolded SNARE motif (ii). Note that the syntaxin-1 molecule was pulled from its C-terminus and I187C to which a DNA handle was attached.
(**B**) Force-extension curves obtained by pulling Munc18-1 bound syntaxin-1 in the absence or presence of 2 µM tomosyn SNARE motif in the solution.
(**C**) Extension-time trajectories at constant forces showing reversible unfolding and refolding of the Munc18-1-bound syntaxin.
Abbreviations: N-terminal regulatory domain (NRD)

**Figure S9: SNARE-truncating constructs support neuronal morphology.**
(**A-D**) Morphological analysis of neurons infected with different mutant constructs (see Figure 5). Control n = 26 – 27/3, cDKO n = 27/3, + WT n = 29/3, + FA n = 33 – 34/3, + ΔPB n = 30 – 31/3, + ΔSNARE n = 36 – 38/3, + WDonly n = 34/3. (**A**) Example images of autaptic hippocampal neurons immunostained for MAP2, synaptophysin-1, and tomosyn-1. Dashed boxes correspond to dendrite zoom-ins in Figure 6F. Scale bar = 50 µm. (**B**) Somatic intensity of tomosyn-1 normalized to controls. (**C**) Total dendrite length as derived from the MAP2 mask. (**D**) Synapse density as derived from the synaptophysin-1-positive puncta within the MAP2 mask.
(**E**) Absolute EPSC amplitudes.
Bonferroni-corrected alpha-levels: *p<.0083, ***p<. Grey asterisks or 'n.s'. show comparison to + WT group. Abbreviations: n.s. (not significant).

**Figure S10: The partial SNARE-hybrid constructs support neuronal morphology.**
(**A-D**) Morphological analysis of neurons infected with different mutant constructs (see Figure 7). (**A**) Example images of autaptic hippocampal neurons immunostained for MAP2, synaptophysin-1, and tomosyn-1. Scale bar = 50 µm. Control n = 22/4, cDKO n = 23/4, + WT n = 24/4, + Core n = 23/4, + LR

n = 25/4. (**B**) Total dendrite length as derived from the MAP2 mask. (**C**) Synapse density as derived from the synaptophysin-1-positive puncta within the MAP2 mask. (**D**) The ratio of the tomosyn-1 signal intensity in synapses to the signal in the soma was calculated to demonstrate synaptic targeting.
(**E**) Absolute EPSC amplitudes.
Bonferroni-corrected alpha-levels: **p<.0025. Grey asterisks or ‘n.s’. show comparison to + WT group. Abbreviations: n.s. (not significant).
