## Supplemental Statistics Table item 1 for "Tomosyns attenuate SNARE assembly and synaptic depression by binding to VAMP2-containing template complexes"

*Supplemental item 1: Overview of statistical analyses per figure. The results of the one-way ANOVA test whether the more complex model (including information on the experimental group observations belong to) is significantly better at capturing the data than the simpler model. Results are reported as F(between groups df, within groups df) = [F-value], p = [p-value]. P-value thresholds = *<0.05; **<0.01;***<0.001. For post-hoc tests, p-value thresholds were adjusted with a Bonferroni correction (α/number of tests).*

| PARAMETER | GROUP | (N/n) | MEAN±SD | OUTL | ANOVA BETWEEN MODELS | POSTHOC ANOVA ON SUBSETS |
|---|---|---|---|---|---|---|
| SUPPLEMENTAL FIGURE 1 | | | | | | |
| Synaptic tom-1 intensity (a.u.) | Control | 2/31 | 1399 ± 334 | 0 | F(1,58) = 231.08, p<0.0001 *** | |
| | cDKO | 2/30 | 424 ± 129 | 0 | | |
| Dendrite length (mm) | Control | 4/62 | 1764 ± 944 | 0 | F(1,114) = 1.1067, p =0.295 | |
| | cDKO | 4/57 | 1605 ± 889 | 0 | | |
| Synapses/ µm | Control | 4/62 | 0.27 ± 0.1 | 1 | F(1,113) = 1.6357, p=0.2035 | |
| | cDKO | 4/57 | 0.28 ± 0.09 | 0 | | |
| Neurite length (mm), DIV7 | Control | 1/13 | 1048 ± 569 | 0 | F(3,60) = 11.54, p<0.0001 *** | WT-DIV7 vs cDKO-DIV7: F(1,25) = 0.09314, p= 0.7628 |
| | cDKO | 1/14 | 1110 ± 471 | 0 | | WT-DIV7 vs WT-DIV15: F(1,30) = 18.26, p= 0.0002*** |
| Neurite length (mm), DIV15 | Control | 1/19 | 2432 ± 1065 | 0 | | WT-DIV15 vs cDKO-DIV15: F(1,35) = 0.8302, p= 0.3685 |
| | cDKO | 1/18 | 2136 ± 899 | 0 | | cDKO-DIV7 vs cDKO-DIV15: F(1,30) = 14.96, p= 0.0005** |
| Synapses/µm, DIV7 | Control | 1/13 | 0.11 ± 0.05 | 0 | F(3,59) = 55.81, p<0.0001 *** | WT-DIV7 vs cDKO-DIV7:F(1,25) = 0.02081, p=0.8865 |
| | cDKO | 1/14 | 0.11 ± 0.05 | 0 | | WT-DIV7 vs WT-DIV15: F(1,29) = 67.93, p<0.0001 *** |
| Synapses/µm, DIV15 | Control | 1/19 | 0.33 ± 0.14 | 1 | | WT-DIV15 vs cDKO-DIV15: F(1,34) = 1.17, p= 0.2869 |
| | cDKO | 1/18 | 0.32 ± 0.07 | 0 | | cDKO-DIV7 vs cDKO-DIV15: F(1,30) = 100.9, p<0.0001 |
| Intensity Tom, DIV7 | Control | 1/13 | 1399 ± 406 | 0 | F(3,58) = 101.7, p<0.0001 *** | WT-DIV7 vs cDKO-DIV7: F(1,24) = 55.84, p<0.0001 *** |
| | cDKO | 1/14 | 624 ± 352 | 1 | | WT-DIV7 vs WT-DIV15: F(1,29) = 1.806, p= 0.1895 |
| Intensity Tom, DIV15 | Control | 1/19 | 1499 ± 268 | 1 | | WT-DIV15 vs cDKO-DIV15: F(1,34) = 381, p<0.0001 *** |
| | cDKO | 1/18 | 412 ± 161 | 0 | | cDKO-DIV7 vs cDKO-DIV15: F(1,29) = 5.912, p= 0.0215 |
| FIGURE 1 + SUPPLEMENTAL FIGURE 2 | | | | | | |
| mEPSC frequency | Control | 6/47 | 6.54 ± 6.54 | 1 | F(1,83) = 111.66, p<0.0001 *** | |
| | cDKO | 6/44 | 27.11 ± 13.87 | 0 | | |
| mEPSC amplitude | Control | 6/47 | 31.16 ± 5.85 | 1 | F(1,83) = 3.0747, p=0.0832 | |
| | cDKO | 6/44 | 32.84 ± 5.68 | 0 | | |
| EPSC amplitude (nA) | Control | 6/50 | 8.4 ± 6.8 | 1 | F(1,91) = 16.435, p=0.0001 *** | |
| | cDKO | 6/49 | 12.2 ± 5.3 | 0 | | |
| EPSC charge (pC) | Control | 6/50 | 124 ± 145 | 1 | F(1,90) = 14.633, p=0.0002 *** | |
| | cDKO | 6/49 | 194 ± 121 | 1 | | |
| EPSC rise time (ms) | Control | 6/44 | 1.68 ± 0.64 | 2 | F(1,83) = 0.1368, p=0.7124 | |
| | cDKO | 6/49 | 0.64 ± 0.47 | 1 | | |
| EPSC decay time (ms) | Control | 6/44 | 6.92 ± 2.81 | 1 | F(1,85) = 3.6273, p=0.0602 | |
| | cDKO | 6/49 | 7.67 ± 2.56 | 0 | | |
| EPSC half-width (ms) | Control | 6/44 | 9.46 ± 2.96 | 1 | F(1,83) = 2.9289, p=0.0907 | |
| | cDKO | 6/49 | 10.22 ± 3.06 | 0 | | |
| PAIRED PULSE RATIO (#2/#1) | | | | | | |
| *20 ms ISI* | Control | 6/45 | 0.95 ± 0.35 | 1 | F(1,83) = 17.249, p<0.0001 *** | |
| | cDKO | 6/47 | 0.71 ± 0.23 | 1 | | |
| *50 ms ISI* | Control | 6/37 | 1.11 ± 0.41 | 1 | F(1,70) = 27.698, p<0.0001*** | |
| | cDKO | 6/42 | 0.74 ± 0.20 | 1 | | |
| *100 ms ISI* | Control | 6/39 | 1.04 ± 0.33 | 0 | F(1,76) = 22.77, p<0.0001*** | |
| | cDKO | 6/44 | 0.75 ± 0.17 | 0 | | |
| *200 ms ISI* | Control | 6/47 | 0.95 ± 0.19 | 1 | F(1,86) = 37.859, p<0.0001*** | |
| | cDKO | 6/47 | 0.74 ± 0.14 | 0 | | |
| SHORT-TERM PLASTICITY (#5/#1) | | | | | | |
| *5 Hz train* | Control | 6/47 | 0.90 ± 0.36 | 1 | F(1,86) = 69.151, p<0.0001*** | |
| | cDKO | 6/47 | 0.50 ± 0.15 | 0 | | |
| *10 Hz train* | Control | 6/39 | 0.85 ± 0.43 | 1 | F(1,75) = 32.539, p<0.0001*** | |
| | cDKO | 6/44 | 0.44 ± 0.16 | 0 | | |
| *20 Hz train* | Control | 6/39 | 0.74 ± 0.37 | 1 | F(1,70) = 35.164, p<0.0001*** | |
| | cDKO | 6/42 | 0.38 ± 0.20 | 1 | | |
| SUPPLEMENTAL FIGURE 3 | | | | | | |
| PPR (#2/#1), 50 ms IPI | | | | | | |
| *1mM [$Ca^{2+}$]e* | Control | 4/13 | 1.57 ± 0.54 | 0 | F(1,22) = 9.6195, p= 0.0052** | |
| | cDKO | 4/14 | 1.12 ± 0.14 | 0 | | |
| *1.5mM [$Ca^{2+}$]e* | Control | 4/9 | 1.18 ± 0.33 | 0 | F(1,18) = 8.7033, p=0.0086** | |
| | cDKO | 4/14 | 0.85 ± 0.16 | 0 | | |
| *2mM [$Ca^{2+}$]e* | Control | 4/13 | 0.98 ± 0.20 | 0 | F(1,22) = 20.336, p= 0.0002*** | |
| | cDKO | 4/14 | 0.72 ± 0.08 | 0 | | |
| *4mM [$Ca^{2+}$]e* | Control | 4/9 | 0.74 ± 0.14 | 0 | F(1,15) = 12.788, p=0.0028** | |
| | cDKO | 4/11 | 0.56 ± 0.07 | 0 | | |
| *8mM [$Ca^{2+}$]e* | Control | 4/9 | 0.63 ± 0.1 | 0 | F(1,14) = 5.9322, p=0.0288* | |
| | cDKO | 4/10 | 0.51 ± 0.08 | 0 | | |
| *Normalized EPSC amp. (to 2mM)* | | | | | | |
| KD $Ca^{2+}$ (mM) | Control | 4/9 | 1.61 ± 0.35 | 0 | F(1,15) = 6.8652, p= 0.0193* | |
| | cDKO | 4/11 | 1.27 ± 0.2 | 0 | | |
| Hill coefficient (*c*) | Control | 4/9 | 3.02 ± 0.57 | 0 | F(1,15) = 0.0815, p=0.7792 | |
| | cDKO | 4/11 | 2.87 ± 0.45 | 0 | | |
| FIGURE 2 + SUPPLEMENTAL FIGURE 4 | | | | | | |
| RRP size (nC) - manual | Control | 6/25 | 2.45 ± 1.26 | 0 | F(1,43) = 1.778, p=0.1894 | |
| | cDKO | 6/26 | 2.15 ± 1.59 | 0 | | |
| Pves (EPSC/RRP) - manual | Control | 6/24 | 0.04 ± 0.03 | 0 | F(1,42) = 6.6264, p=0.0137* | |
| | cDKO | 6/25 | 0.07 ± 0.06 | 0 | | |
| 250/500mM - manual | Control | 6/25 | 0.45 ± 0.21 | 0 | F(1,43) = 42.129, p<0.0001*** | |
| | cDKO | 6/26 | 0.85 ± 0.24 | 0 | | |
| Refilled pool (% of initial) | Control | 6/19 | 0.79 ± 0.07 | 0 | F(1,32) = 0.2302, p= 0.6347 | |
| | cDKO | 6/20 | 0.80 ± 0.09 | 0 | | |

| | | | | | | |
|---|---|---|---|---|---|---|
| RRP size (nC) – fitted | Control | 6/16 | 1.99 ± 1.02 | 0 | F(1,27) = 0.6872, p=0.4144 | |
| | cDKO | 6/18 | 1.33 ± 0.91 | 0 | | |
| Pves (EPSC/RRP) – fitted | Control | 6/15 | 0.05 ± 0.04 | 0 | F(1,23) = 2.0802, p=0.1627 | |
| | cDKO | 6/15 | 0.10 ± 0.06 | 0 | | |
| 250/500mM - fitted | Control | 6/10 | 0.55 ± 0.09 | 0 | F(1,19) = 129.29, p<0.0001*** | |
| | cDKO | 6/15 | 0.92 ± 0.06 | 0 | | |
| $K_{2,max}$($s^{-1}$) | | | | | | |
| *0mM HS* | Control | 6/13 | 0.0001 ± 0.00008 | 0 | F(1,14) = 6.3655, p= 0.02436* | |
| | cDKO | 6/7 | 0.0013 ± 0.0010 | 0 | | |
| *250mM HS* | Control | 6/12 | 0.43 ± 0.81 | 0 | F(1,22) = 21.344, p=0.0001*** | |
| | cDKO | 6/17 | 0.80 ± 0.42 | 1 | | |
| *500mM HS* | Control | 6/16 | 3.02 ± 3.53 | 0 | F(1,26) = 41.962, p<0.0001*** | |
| | cDKO | 6/18 | 5.13 ± 1.19 | 0 | | |
| $\Delta E_a$(RT) | | | | | | |
| *0mM HS* | Control | 6/13 | 0.00 ± 0.98 | 0 | F(1,14) = 21.025, p= 0.0004*** | |
| | cDKO | 6/7 | -2.43 ± 0.68 | 0 | | |
| *250mM HS* | Control | 6/12 | -7.84 ± 0.85 | 0 | F(1,23) = 21.363, p= 0.0001*** | |
| | cDKO | 6/17 | -8.95 ± 0.44 | 0 | | |
| *500mM HS* | Control | 6/16 | -10.1 ± 0.63 | 0 | F(1,26) = 48.629, p< 0.0001*** | |
| | cDKO | 6/18 | -10.9 ± 0.24 | 1 | | |
| *Priming rate* $K_1$($s^{-1}$) | Control | 6/16 | 0.17 ± 0.20 | 1 | F(1,24) = 3.7675, p=0.06409 | |
| | cDKO | 6/16 | 0.07 ± 0.14 | 0 | | |
| *Unpriming rate* $K_{-1}$($s^{-}$ | Control | 6/16 | 0.07 ± 0.07 | 1 | F(1,24) = 3.9587, p=0.05814 | |
| | cDKO | 6/16 | 0.04 ± 0.06 | 0 | | |
| SYNAPTIC RECOVERY AFTER TRAIN STIMULATION | | | | | | |
| *40Hz train* | | | | | | |
| Norm. EPSC – R1 | Control | 6/33 | 1.00 ± 0.31 | 0 | F(1,64) = 53.489, p<0.0001*** | |
| | cDKO | 6/38 | 0.59 ± 0.13 | 0 | | |
| Norm. EPSC – R2 | Control | 6/31 | 0.95 ± 0.21 | 0 | F(1,59) = 1.7315, p=0.1933 | |
| | cDKO | 6/35 | 0.90 ± 0.14 | 0 | | |
| $1^{st}$ EPSC (nA) | Control | 6/33 | 6.95 ± 4.85 | 1 | F(1,62) = 7.5857, p<0.007711** | |
| | cDKO | 6/38 | 9.51 ± 5.08 | 1 | | |
| R1 EPSC (nA) | Control | 6/33 | 6.38 ± 4.42 | 1 | F(1,62) = 0.4848, p=0.4889 | |
| | cDKO | 6/38 | 5.57 ± 2.94 | 1 | | |
| R2 EPSC (nA) | Control | 6/31 | 6.36 ± 4.89 | 1 | F(1,57) = 4.8583, p=0.03157* | |
| | cDKO | 6/35 | 8.14 ± 4.54 | 1 | | |
| RRP size (nC) | Control | 6/33 | 0.78 ± 0.51 | 0 | F(1,63) = 2.6424, p=0.109 | |
| | cDKO | 6/38 | 0.61 ± 0.46 | 1 | | |
| Refilling rate (nC/s) | Control | 6/33 | 1.05 ± 1.00 | 1 | F(1,61) = 2.2141, p=0.1419 | |
| | cDKO | 6/38 | 0.83 ± 0.61 | 2 | | |
| FITTED PARAMETERS RECOVERY 500mM HS | | | | | | |
| $K_{2,max}$($s^{-1}$) | Control | 6/12 | 2.06 ± 1.31 | 0 | F(1,16) = 0.738, p= 0.403 | |
| | cDKO | 6/11 | 2.52 ± 1.10 | 0 | | |
| $\Delta E_a$(RT) | Control | 6/12 | -9.83 ± 0.64 | 0 | F(1,16) = 0.6905, p=0.4182 | |
| | cDKO | 6/11 | -10.03 ± 0.72 | 0 | | |
| SUPPLEMENTAL FIGURE 5 | | | | | | |
| Docked SVs | Control | 4/132 | 8.11 ± 3.78 | 1 | L ratio (2,339) = 7.152772, p=0.028 * | Control-cDKO: p=0.0424 |
| | cDKO | 4/89 | 6.57 ± 3.45 | 1 | | Control-WT: p=0.0153* |
| | +WT | 4/128 | 6.87 ± 4.57 | 2 | | cDKO-WT: p=0.804 |
| AZ length (μm) | Control | 4/132 | 623 ± 271 | 1 | L ratio (2,335) = 3.175503, p=0.2044 | |
| | cDKO | 4/89 | 580 ± 308 | 4 | | |
| | +WT | 4/128 | 643 ± 285 | 3 | | |
| Total SV number | Control | 4/132 | 154 ± 96 | 3 | L ratio (2,335) = 2.343999, p=0.3097 | |
| | cDKO | 4/89 | 134 ± 92 | 3 | | |
| | +WT | 4/128 | 143 ± 99 | 2 | | |
| FIGURE 3 + SUPPLEMENTAL FIGURE 6 | | | | | | |
| EPSC amp. (nA) | cDKO | 4/17 | 9.73 ± 5.33 | 0 | F(2,52) = 7.5077, p=0.001366** | *vsWT:* F(1,32) = 12.3, p=0.001366** |
| | +WT | 4/21 | 5.03 ± 6.13 | 1 | | |
| | +Hybrid | 4/21 | 10.75 ± 7.33 | 0 | | *vsWT:* F(1,36) = 11.866, p=0.001468** |
| STP ($5^{th}/1^{st}$) | cDKO | 4/17 | 0.35 ± 0.15 | 0 | F(2,53) = 15.079, p<0.0001*** | *vsWT:* F(1,33) = 25.495, p<0.0001*** |
| | +WT | 4/21 | 0.93 ± 0.47 | 0 | | |
| | +Hybrid | 4/21 | 0.54 ± 0.27 | 0 | | *vsWT:* F(1,37) = 10.764, p=0.002262** |
| EPSC recovery | cDKO | 4/14 | 0.51 ± 0.10 | 0 | F(2,46) = 28.714, p<0.0001*** | *vsWT:* F(1,29) = 36.203, p<0.0001*** |
| | +WT | 4/20 | 1.48 ± 0.67 | 0 | | |
| | +Hybrid | 4/18 | 0.65 ± 0.23 | 0 | | *vsWT:* F(1,33) = 30.101, p<0.0001*** |
| Norm. synaptic tom-1 | Control | 3/35 | 1.00 ± 0.26 | 0 | F(3,125) = 50.516, p<0.0001*** | *vsWT:* F(1,63) = 0.7486, p=0.3902 |
| | cDKO | 3/30 | 0.35 ± 0.16 | 0 | | *vsWT:* F(1,58) = 50.516, p<0.0001*** |
| | +WT | 3/33 | 1.12 ± 0.48 | 1 | | |
| | +hybrid | 3/35 | 1.69 ± 0.88 | 1 | | *vsWT:* F(1,62) = 15.887, p=0.0001797*** |
| Synaptic / somatic tomosyn | Control | 3/35 | 0.66 ± 0.40 | 0 | F(3,124) = 22.94, p<0.0001*** | *vsWT:* F(1,63) = 1.1227, p=0.2934 |
| | cDKO | 3/30 | 1.04 ± 0.44 | 0 | | *vsWT:* F(1,58) = 45.35, p<0.0001*** |
| | +WT | 3/33 | 0.60 ± 0.25 | 1 | | |
| | +hybrid | 3/35 | 0.65 ± 0.27 | 1 | | *vsWT:* F(1,63) = 0.8051, p=0.373 |
| Total dendrite length (mm) | Control | 3/35 | 2.36 ± 1.07 | 0 | F(3,127) = 6.0089, p=0.0007*** | *vsWT:* F(1,64) = 3.6022, p=0.06222 |
| | cDKO | 3/30 | 1.72 ± 0.69 | 0 | | *vsWT:* F(1,59) = 3.118, p= 0.0826 |
| | +WT | 3/33 | 2.05 ± 0.87 | 0 | | |
| | +hybrid | 3/35 | 2.45 ± 1.17 | 0 | | *vsWT:* F(1,64) = 4.6172, p= 0.03544 |
| Synapses/μm dendrite | Control | 3/35 | 0.28 ± 0.09 | 0 | F(3,127) = 1.237, p= 0.2991 | |
| | cDKO | 3/30 | 0.26 ± 0.08 | 0 | | |
| | +WT | 3/33 | 0.30 ± 0.09 | 0 | | |
| | +hybrid | 3/35 | 0.30 ± 0.10 | 0 | | |
| FIGURE 6 + SUPPLEMENTAL FIGURE 9 | | | | | | |
| EPSC amp. (nA) | Control | 6/21 | 7.23 ± 4.90 | 0 | F(6,106) = 2.0689, p=0.06294 | |
| | cDKO | 8/25 | 9.89 ± 6.77 | 1 | | |

| | | | | | | |
|---|---|---|---|---|---|---|
| | +WT | 6/27 | 4.69 ± 4.22 | 1 | | |
| | +FA | 5/18 | 8.81 ± 6.61 | 0 | | |
| | +ΔPB | 4/13 | 9.20 ± 5.68 | 0 | | |
| | +ΔSNARE | 4/8 | 7.08 ± 5.46 | 0 | | |
| | +WDonly | 3/10 | 8.07 ± 4.83 | 0 | | |
| STP ($5^{th}/1^{st}$) | Control | 6/21 | 0.78 ± 0.41 | 0 | F(6,108) = 13.21, p<0.0001 *** | *vsKO:* F(1,37) = 19.352, p=0.00011*** |
| | cDKO | 8/25 | 0.44 ± 0.16 | 0 | | |
| | +WT | 6/27 | 1.25 ± 0.72 | 0 | | *vsKO:* F(1,43) = 26.297, p<0.0001*** |
| | +FA | 5/18 | 0.50 ± 0.14 | 0 | | *vsKO:* F(1,34) = 1.1017, p=0.3013 |
| | +ΔPB | 4/13 | 0.37 ± 0.08 | 0 | | *vsKO:* F(1,29) = 0.0239, p= 0.8781 |
| | +ΔSNARE | 4/8 | 0.44 ± 0.20 | 0 | | *vsKO:* F(1,24) = 0.0782, p= 0.7821 |
| | +WDonly | 3/10 | 0.41 ± 0.16 | 0 | | *vsKO:* F(1,26) = 1.3305, p=0.2592 |
| EPSC recovery | Control | 6/21 | 0.94 ± 0.36 | 0 | F(6,96) = 16.175, p<0.0001 *** | *vsKO:* F(1,34) = 30.275, p=0.00011*** |
| | cDKO | 8/25 | 0.53 ± 0.16 | 0 | | |
| | +WT | 6/25 | 1.49 ± 0.79 | 1 | | vsKO: F(1,37) = 41.166, p<0.0001*** |
| | +FA | 5/16 | 0.69 ± 0.24 | 0 | | *vsKO:* F(1,29) = 4.7929, p=0.03677 |
| | +ΔPB | 4/12 | 0.52 ± 0.15 | 0 | | *vsKO:* F(1,25) = 0.5245, p=0.4757 |
| | +ΔSNARE | 3/6 | 0.56 ± 0.15 | 0 | | *vsKO:* F(1,19) = 0.1732, p=0.682 |
| | +WDonly | 2/9 | 0.52 ± 0.14 | 0 | | *vsKO:* F(1,22) = 0.0363, p=0.8506 |
| Norm. synaptic tom-1 levels | Control | 3/27 | 1.00 ± 0.29 | 1 | F(6,206) = 25.823, p<0.0001 *** | *vsWT:* F(1,51) = 1.9089, p=0.1731 |
| | cDKO | 3/27 | 0.22 ± 0.05 | 0 | | |
| | +WT | 3/29 | 1.13 ± 0.85 | 0 | | vsKO: F(1,52) = 48.473, p<0.0001*** |
| | +FA | 3/34 | 0.56 ± 0.40 | 1 | | *vsWT:* F(1,58) = 19.222, p<0.0001*** |
| | +ΔPB | 3/31 | 1.48 ± 1.03 | 0 | | *vsWT:* F(1,56) = 4.1628, p=0.04605 |
| | +ΔSNARE | 3/38 | 0.42 ± 0.21 | 1 | | *vsWT:* F(1,61) = 29.97, p<0.0001*** |
| | +WDonly | 3/34 | 0.64 ± 0.30 | 1 | | *vsWT:* F(1,58) = 13.402, p= 0.00054** |
| Norm. somatic tom-1 levels | Control | 3/26 | 1.00 ± 0.18 | 1 | F(6,205) = 34.025, p<0.0001 *** | *vsWT:* F(1,50) = 2.4075, p=0.1271 |
| | cDKO | 3/27 | 0.26 ± 0.11 | 0 | | vsWT: F(1,52) = 78.04, p<0.0001*** |
| | +WT | 3/29 | 0.90 ± 0.41 | 0 | | |
| | +FA | 3/33 | 0.66 ± 0.38 | 0 | | *vsWT:* F(1,58) = 7.247, p=0.009266 |
| | +ΔPB | 3/30 | 1.09 ± 0.22 | 0 | | *vsWT:* F(1,55) = 8.589, p= 0.004917* |
| | +ΔSNARE | 3/36 | 0.68 ± 0.25 | 0 | | *vsWT:* F(1,61) = 7.1465, p=0.009623 |
| | +WDonly | 3/34 | 0.96 ± 0.27 | 0 | | *vsWT:* F(1,59) = 1.7049, p= 0.1967 |
| Total dendritic length (mm) | Control | 3/27 | 1.00 ± 0.45 | 0 | F(6,210) = 2.8711, p= 0.0104* | *vsWT:* F(1,52) = 3.7978, p= 0.05672 |
| | cDKO | 3/27 | 1.11 ± 0.62 | 0 | | *vsWT:* F(1,52) = 1.3874, p= 0.2442 |
| | +WT | 3/29 | 1.29 ± 0.79 | 0 | | |
| | +FA | 3/34 | 1.45 ± 0.89 | 1 | | *vsWT:* F(1,58) = 0.1994, p= 0.6569 |
| | +ΔPB | 3/31 | 1.58 ± 0.93 | 0 | | *vsWT:* F(1,56) = 1.9107, p= 0.1724 |
| | +ΔSNARE | 3/38 | 1.42 ± 0.66 | 0 | | *vsWT:* F(1,63) = 1.2786, p= 0.2624 |
| | +WDonly | 3/34 | 1.44 ± 0.75 | 0 | | *vsWT:* F(1,59) = 1.2978, p= 0.2592 |
| Synapses/µm dendrite | Control | 3/27 | 0.26 ± 0.07 | 0 | F(6,211) = 0.5898, p= 0.7383 | |
| | cDKO | 3/27 | 0.25 ± 0.08 | 0 | | |
| | +WT | 3/29 | 0.26 ± 0.07 | 0 | | |
| | +FA | 3/34 | 0.24 ± 0.09 | 0 | | |
| | +ΔPB | 3/31 | 0.23 ± 0.06 | 0 | | |
| | +ΔSNARE | 3/38 | 0.25 ± 0.06 | 0 | | |
| | +WDonly | 3/34 | 0.24 ± 0.07 | 0 | | |
| FIGURE 7 + SUPPLEMENTAL FIGURE S10 | | | | | | |
| EPSC amp. (nA) | cDKO | 4/20 | 11.30 ± 5.99 | 0 | F(3,81) = 4.542, p=0.005406** | *vsWT:* F(1,37) = 8.8633, p=0.005108* |
| | +WT | 4/22 | 6.78 ± 4.89 | 0 | | |
| | +Core | 4/24 | 7.44 ± 7.74 | 1 | | *vsWT:* F(1,40) = 0.0058, p=0.9399 |
| | +LR | 4/23 | 7.95 ± 5.59 | 0 | | *vsWT:* F(1,40) = 1.059, p=0.3096 |
| STP ($5^{th}/1^{st}$) | cDKO | 4/20 | 0.35 ± 0.09 | 0 | F(3,81) = 28.328, p<0.0001*** | *vsWT:* F(1,37) = 52.416, p<0.0001*** |
| | +WT | 4/22 | 0.97 ± 0.44 | 0 | | |
| | +Core | 4/24 | 0.56 ± 0.17 | 0 | | *vsWT:* F(1,41) = 34.323, p<0.0001*** |
| | +LR | 4/23 | 0.58 ± 0.41 | 1 | | *vsWT:* F(1,39) = 29.404, p<0.0001*** |
| EPSC recovery | cDKO | 4/19 | 0.54 ± 0.11 | 0 | F(3,77) = 15.46, p<0.0001*** | *vsWT:* F(1,35) = 36.875, p<0.0001*** |
| | +WT | 4/22 | 1.52 ± 0.98 | 1 | | |
| | +Core | 4/22 | 1.00 ± 0.62 | 0 | | *vsWT:* F(1,38) = 9.1997, p=0.004347* |
| | +LR | 4/22 | 0.79 ± 0.33 | 0 | | *vsWT:* F(1,38) = 21.067, p<0.0001*** |
| Norm. synaptic tom-1 levels | Control | 4/22 | 1.00 ± 0.28 | 0 | F(4,109) = 16.672, p<0.0001*** | *vsWT:* F(1,41) = 0.0987, p=0.7549 |
| | cDKO | 4/23 | 0.30 ± 0.12 | 0 | | *vsWT:* F(1,42) = 57.035, p<0.0001*** |
| | +WT | 4/24 | 1.04 ± 0.52 | 0 | | |
| | +Core | 4/23 | 0.91 ± 0.58 | 0 | | *vsWT:* F(1,42) = 0.9359, p=0.3389 |
| | +LR | 4/25 | 1.21 ± 0.53 | 0 | | *vsWT:* F(1,44) = 0.9231, p=0.3419 |
| Synaptic / somatic tomosyn | Control | 4/22 | 0.83 ± 0.40 | 0 | F(4,106) = 3.3321, p=0.013* | *vsWT:* F(1,40) = 1.8309, p=0.1836 |
| | cDKO | 4/23 | 0.98 ± 0.37 | 0 | | *vsWT:* F(1,40) = 16.131, p=0.0002531** |
| | +WT | 4/24 | 0.74 ± 0.28 | 1 | | |
| | +Core | 4/23 | 0.74 ± 0.33 | 0 | | *vsWT:* F(1,41) = 0.2538, p=0.6171 |
| | +LR | 4/25 | 0.82 ± 0.27 | 0 | | *vsWT:* F(1,42) = 4.6526, p=0.03678 |
| Total dendritic length (mm) | Control | 4/22 | 1.72 ± 0.53 | 0 | F(4,109) = 2.3891, p=0.0553 | *vsWT:* F(1,41) = 0.4295, p=0.5159 |
| | cDKO | 4/23 | 1.56 ± 0.57 | 0 | | *vsWT:* F(1,42) = 2.2197, p= 0.1437 |
| | +WT | 4/24 | 1.86 ± 0.85 | 0 | | |
| | +Core | 4/23 | 2.12 ± 1.05 | 0 | | *vsWT:* F(1,42) = 1.4927, p= 0.2286 |
| | +LR | 4/25 | 2.11 ± 0.91 | 0 | | *vsWT:* F(1,44) = 0.9108, p= 0.3451 |
| Synapses/µm dendrite | Control | 4/22 | 0.24 ± 0.07 | 0 | F(4,109) = 0.8741, p= 0.482 | |
| | cDKO | 4/23 | 0.23 ± 0.10 | 0 | | |
| | +WT | 4/24 | 0.21 ± 0.07 | 0 | | |
| | +Core | 4/23 | 0.22 ± 0.07 | 0 | | |
| | +LR | 4/25 | 0.24 ± 0.07 | 0 | | |
